# Identification of a human hematopoietic stem cell subset that retains memory of inflammatory stress

**DOI:** 10.1101/2023.09.11.557271

**Authors:** Andy G.X. Zeng, Murtaza S. Nagree, Niels Asger Jakobsen, Sayyam Shah, Alex Murison, Jin-Gyu Cheong, Sven Turkalj, Isabel N.X. Lim, Liqing Jin, Joana Araújo, Alicia G. Aguilar-Navarro, Darrien Parris, Jessica McLeod, Hyerin Kim, Ho Seok Lee, Lin Zhang, Mason Boulanger, Elvin Wagenblast, Eugenia Flores-Figueroa, Bo Wang, Gregory W. Schwartz, Leonard D. Shultz, Steven Z. Josefowicz, Paresh Vyas, John E. Dick, Stephanie Z. Xie

**Author notes:** These authors contributed equally (Andy G.X. Zeng, Murtaza S. Nagree, Niels Asger Jakobsen). Co-senior authors (Paresh Vyas, John E. Dick, Stephanie Z. Xie).

## Abstract

Inflammation activates many blood cell types, driving aging and malignancy. Yet, hematopoietic stem cells (HSCs) survive a lifetime of infection to sustain life-long blood production. To understand HSC adaptation to inflammation, we developed xenograft inflammation-recovery models and performed single cell multiomics on isolated human HSC. Two transcriptionally and epigenetically distinct HSC subsets expressing canonical HSC programs were identified. Only one showed sustained transcriptional and epigenetic changes after recovery from inflammatory treatments. This HSC inflammatory memory (HSC-iM) program is enriched in memory T cells and HSCs from recovered COVID-19 patients. Importantly, HSC-iM accumulates with age and with clonal hematopoiesis. Overall, heritable molecular alterations in a subset of human HSCs, an adaptation to long-term inflammatory stress, may predispose to heightened age-related risk of blood cancer and infection.

**One-Sentence Summary:** Inflammation across a lifetime rewires human HSCs to produce a distinct HSC subset with both beneficial and deleterious fitness consequences.

## Introduction

Humans have an enormous demand (∼10^11^ cells daily) for hematopoietic output (*1*). Meeting this need over a lifetime is achieved by a complex cellular hierarchy with a heterogeneous pool of dormant hematopoietic stem cells (HSCs) at its apex. Mature blood cells with a finite life span are continuously replenished by bone marrow (BM) hematopoietic stem and progenitor cells (HSPCs) producing three million cells per second in human adults, via a tightly controlled process (*2*). Adult humans are estimated to have 50,000 to 200,000 HSCs contributing to hematopoiesis at any one time (*3*). HSCs are distinguished from their immediate downstream progenitors (collectively termed HSPCs) by their cycling status and capacity for self-renewal (*4*). HSCs show functional decline with age (*5–10*), with a variable but often marked decrease in HSC clonal diversity (*11*), and a consequent increase in incidence of clonal hematopoiesis (CH) (*12*) and risk of blood malignancy (*13*). The coordination between daily hematopoietic output and maintenance of the HSC pool over the lifetime of an individual is not well-understood, especially in the context of inflammatory stress, such as repeated infections, which induce signaling programs known to activate HSCs (*14–17*). Indeed, the transitions HSCs undergo when exiting quiescence towards activation are essential for human HSC function (*17–21*). Such transitions become dysregulated in human HSCs with aging, and inflammatory stress, but are poorly understood both from a cellular and molecular standpoint (*18, 19, 22*). Additionally, HSCs are not homogeneous. We previously identified transcriptional, epigenetic, and functional heterogeneity in human HSCs, with inflammatory pathways being one of the most prominent sources of such heterogeneity (*15, 17, 20, 21, 23, 24*). Together, these data imply that cell type-specific responses to inflammation within heterogeneous HSC subsets might be the driver that maintains long-term homeostasis of the HSC pool in the face of a lifetime of inflammatory insults, (*25–27*).

Human lineage tracking data show pre-leukemic CH mutations often arise many decades before disease onset (*28–31*). Importantly, small clones bearing CH mutations are found almost ubiquitously in 50-60-year old adults (*32*), but only a small proportion expand to reach a readily detectable clone size. Crucially, the mechanisms regulating CH clone size are poorly understood. In mouse models, acute inflammation activates HSCs, skews them towards myeloid differentiation, and impairs their self-renewal (*16, 33, 34*). Some of these inflammatory response programs include activation of target genes downstream of TNFα via NFkB that regulate HSC survival in both mouse and human settings (*15, 16*). Repeated inflammatory challenges promote sustained epigenetic changes (*35*) and accelerate aging of murine HSCs (*14, 36*). Moreover, inflammation and aging in mice promote expansion of clones bearing CH-associated mutations in *Dnmt3a* and *Tet2* (*37–41*). However, a distinct cellular compartment responding to inflammation or CH-associated mutations within the HSC pool has not previously been described. Additionally, how inflammatory stress, and age-related decline of HSC fitness, is linked to perturbed regulation of dormancy/activation, and to selection of specific HSC clones in CH has not been resolved in humans (*13, 36*). We set out to address these questions. Here, we identify a previously unrecognized human HSC subset that retains memory of prior inflammatory stress, whose transcriptomic and epigenetic signatures are tightly correlated to HSC from aged individuals, from patients who recovered from severe COVID-19 infection, and from individuals with CH.

## Results

### Heterogeneous transcriptional priming of inflammatory response in unperturbed LT-HSCs

Human HSC quiescence-activation status ranges from deeply quiescent to being primed for cell cycle entry, and is intimately tied to variations in regeneration kinetics, 3D chromatin architecture and endolysosomal activity (*15, 17, 21, 23*). To explore how quiescence status correlates with transcriptional inflammatory response programs in unperturbed LT-HSCs, we performed single-cell (sc) RNA-sequencing (scRNA-seq) on highly purified umbilical cord blood (CB) Lin-CD34+CD38-CD45RA-CD90+CD49f+ long-term HSCs (LT-HSCs). Consensus non-negative matrix factorization (cNMF) (*42*) was used to infer gene expression programs that vary across 3,381 LT-HSC transcriptomes (Fig. 1A). Distinct gene expression programs were identified relating to quiescence (Fig. 1, B and C) and inflammatory signaling (Fig. 1, D and E, and Table S1 and S2). To validate these findings, simultaneous single-nucleus gene expression and chromatin accessibility profiling (hereafter scMultiome) was performed on CB HSPCs (n=2 females, n=2 males). We projected the sc-transcriptomes against a hematopoietic reference map (github.com/andygxzeng/BoneMarrowMap; hereafter BoneMarrowMap). In this reference map, transcriptionally-defined HSCs are highly concordant with immunophenotypic LT-HSCs (fig. S1, A and B) and enabled us to identify 15,590 CB HSCs and multipotent progenitors (MPPs) *in silico* from our scMultiome dataset (fig. S1, C). cNMF of their transcriptomes recovered similar gene expression programs corresponding to quiescence, cell cycle priming, myeloid-lymphoid (MyLy) and megakaryocyte-erythroid (MkEry) lineage priming programs, and inflammatory signaling providing independent validation of the scRNA-seq results (fig. S1, D to O, and Table S2 and S3). Notably, 60 recurrent genes were identified that are shared between the top 200 genes driving the two independent inflammatory signaling programs from the two HSC datasets (fig. S1, P to R, and Table S4). Further, chromatin accessibility information from the HSC scMultiome dataset demonstrated that the inflammatory gene expression program was associated with motif accessibility of AP-1 and NF-kB transcription factors (TFs) (fig. S1, S to U). These data, together with previous findings (*17, 21, 23, 24*), demonstrate variable levels of priming for inflammatory response within individual HSC/MPP from inflammation-naive CB.

**Fig. 1.**
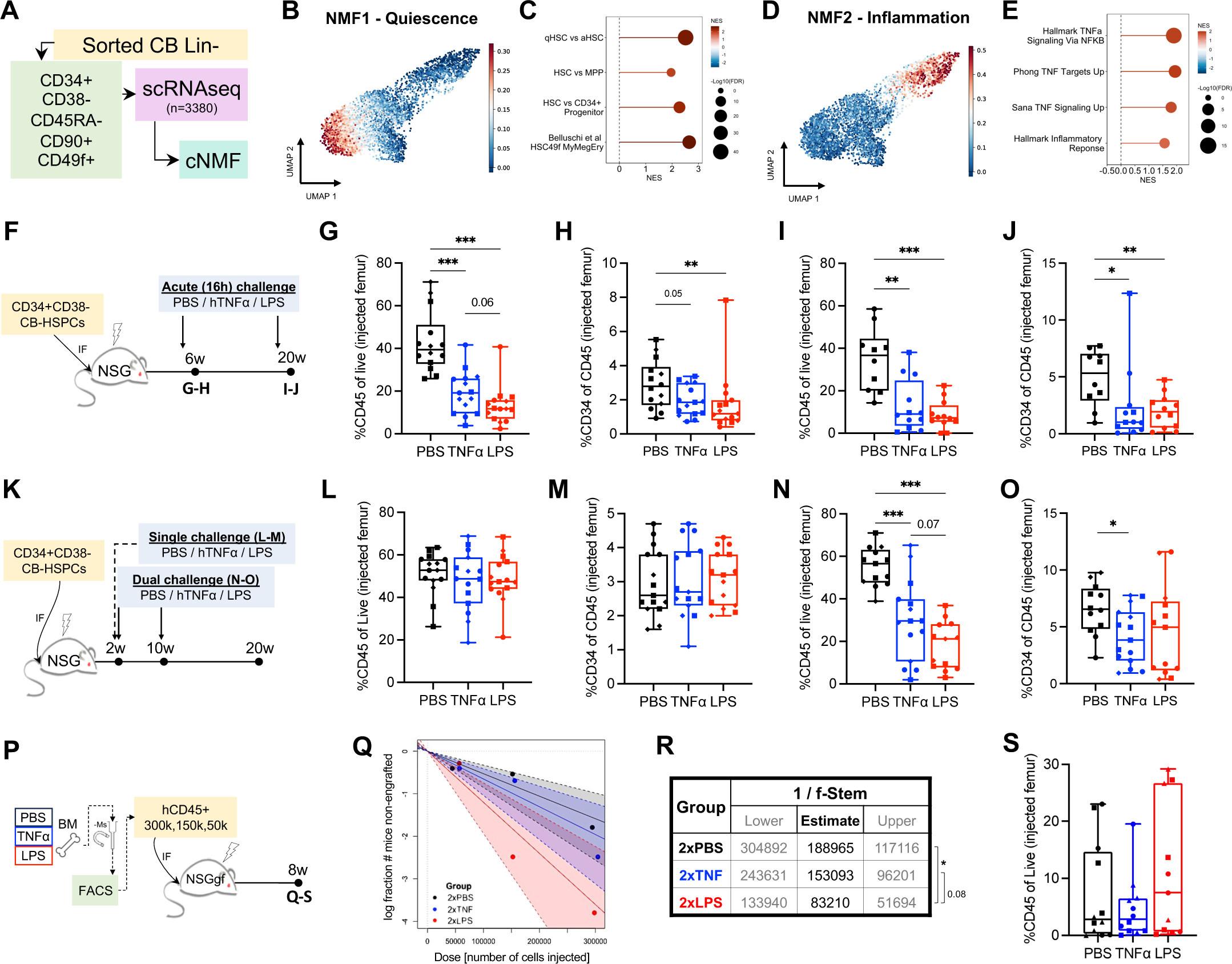
Modeling inflammatory response and recovery in human LT-HSC. **(A)** LT-HSC were purified from umbilical cord blood (CB) and subjected to single-cell (sc) RNA sequencing. Consensus non-negative matrix factorization (cNMF) was performed on 3,381 cells. **(B-C)** Localized enrichment (B) of a cNMF transcriptional program of quiescence, as defined by GSEA enrichment (C). **(D-E)** Localized enrichment of a cNMF transcriptional program of Inflammation as defined by GSEA enrichment (E). **(F-J)** NSG mice xenografted with CB-derived CD34+CD38-cells via intrafemoral (IF) injection after irradiation were challenged with hTNFα or lipopolysaccharide (LPS) at either 6w (G-H) or 20w (I-J) post-transplant. The effect on total human engraftment (G, I) and progenitor compartment (H, J) 16h after inflammatory challenge was measured and compared to mice challenged with vehicle control (PBS). **(K)** Schematic for xenograft model of recovery from single (2w) or dual (2w and 10w) acute inflammatory challenge. **(L-M)** Human engraftment (L) and progenitor composition (M) 18w after a single inflammatory challenge. **(N-O)** Human engraftment (N) and progenitor composition (O) 10w after dual inflammatory challenge. **(P)** Experimental schematic for secondary transplantation with limiting dilution of FACS purified human leukocytes from dual challenged xenografts. **(Q-R)** Stem cell frequency estimates for (P). **(S)** Human engraftment in cohorts from (P) transplanted with 300,000 hCD45+ cells. All engraftment data is presented as box plots with a line at the median, boxes showing quartiles and error bars showing range. Individual points show data for each animal used in the study (n=10-15), and different symbols reflect different CB pools (n=2-3). Data were compared using pairwise Mann-Whitney tests; stem cell frequency estimates were compared using chi-squared tests, *** *p*<0.001, ** *p*<0.01, * *p*<0.05, and *p*<0.1 is shown numerically, while *p*>0.1 is not shown.

### *In vivo* modeling of inflammatory insult and recovery of human HSCs

To determine whether the observed molecular variation in inflammatory response priming is linked to functional variation in human HSCs, we developed a novel xenotransplantation model of inflammation. We focused on the impact of lipopolysaccharide (LPS), which mimics sepsis (*43*), and tumor necrosis factor α (TNFα) which has been associated with deleterious effects on human health in the context of advanced age, COVID-19 and sepsis (*44–47*). Although TNFα signaling has been shown to regulate HSC proliferation and survival (*15, 16, 19, 48*)*, in vivo* inflammatory stress responses in human HSCs following TNFα administration have not been investigated. We first examined whether human TNFα (hTNFα) or LPS could provoke acute inflammatory responses in a human xenograft (Fig. 1F). Mice were treated with PBS, hTNFα, or LPS at two timepoints post-CB transplantation: 6 weeks (w), when the engrafted human HSCs are mostly cycling (Fig. 1, G and H and fig. S2, B and C) and 20w when engrafted HSC are mainly quiescent (Fig. 1, I and J, and fig. S2, D and E) (*24*). Human CD45+ cell abundance and the proportion of CD34+ cells were measured in the injected femur and non-injected femur. Human engraftment within the injected BM derives from a combination of HSCs and progenitors, while the non-injected BM compartment arises from self-renewing HSC (*17*). Both total human CD45+ cells and the proportion of human CD34+ progenitors were reduced in the injected femurs 16 hours after hTNFα/LPS treatment at both 6w (Fig. 1, G and H) and 20w (Fig. 1, I and J) post-transplant compared to PBS controls. The non-injected BM showed more variable response to acute inflammation at 6w vs 20w (fig. S2, A to E). We observed no alterations in the balance between B-lymphoid and myeloid lineages (fig. S3, A to E). Overall, these findings demonstrate a negative impact of acute inflammatory stimuli on human hematopoietic output in a xenograft model, consistent with previous studies of mouse HSCs (*36*).

Next, we tested whether CB xenografts can recover from acute inflammatory insults. A single challenge was performed with PBS, hTNFα, or LPS at 2w post-transplant, and human engraftment patterns subsequently analyzed at 20w post-transplant (Fig. 1K). Overall engraftment in the PBS-injected controls compared to LPS- or hTNFα-treated mice was similar in the injected femurs (Fig. 1, L and M) and non-injected femurs (fig. S2, F to H), demonstrating that human grafts are capable of recovering from a single inflammatory insult, congruent with findings in mice (*36*). No significant differences in lineage distribution were observed (fig. S3, F to H). Furthermore, limiting dilution assays (LDA) into secondary NSG-SGM3 mice did not reveal statistically significant differences in stem cell frequency between mice treated with a single inflammatory challenge of PBS, hTNFα, or LPS (fig. S2, I to K).

To investigate the impact of repeated inflammatory insults on human HSC function, we developed a dual challenge model wherein xenografted NSG mice were treated with PBS, hTNFα, or LPS at 2w and 10w post-transplant followed by a further 10w recovery period (Fig. 1K). The human CD45+ graft in the injected femur at 20w post-transplant remained significantly lower in mice challenged with either LPS or hTNFα compared to PBS despite the recovery period (Fig. 1N). Human CD34+ proportion was also reduced with hTNFα challenge (Fig 1O). Variable engraftment was observed at distal non-injected BM following repeated inflammatory challenge, where only LPS challenge impacted engraftment, possibly reflective of a role for the degree of inflammatory insult (fig. S2, L and M). The effect of dual challenge on lineage distribution after recovery of the human grafts was variable (fig. S2N, and fig. S3, I and J). An increase in the proportion of CD3+ T cells was observed upon recovery from LPS or hTNFα at the expense of CD19+ B cells (fig. S3, I and J). CapTCR-seq showed that this was not due to clonal T cell expansion (fig. S3, K to M). To study whether a radiation-altered microenvironment may have impacted the human HSC response to dual inflammatory challenge, we adapted our model to NSGW41 recipient mice (fig. S4A). Endogenous murine HSCs in NSGW41 mice are impaired due to a Kit deficiency, allowing human HSC engraftment without irradiation (*49*). Robust human CD45+ leukocytic and GlyA+ erythrocytic grafts were obtained 20w post-transplant in PBS-treated mice, and grafts were significantly reduced after recovery from dual LPS or hTNFα challenge (fig. S4, B to G). No significant changes in the proportion of CD34+ cells or mature lineages were observed (fig. S4, H to K). These findings with NSGW41 recipients argue that an irradiated niche did not play a role in the response to inflammatory treatment.

Despite the impaired hematopoietic output observed at 20w following dual inflammatory challenge, secondary transplant with LDA (Fig. 1P) demonstrated that HSC with serial engraftment capacity were still present in recipient BM at similar frequency comparing PBS to hTNFα groups, and with modest increase for the LPS group (Fig. 1, Q to S). This is in contrast to mouse models of chronic inflammation in which HSCs failed to recover functional potency up to a year after inflammatory challenge (*14, 36*). Thus, our model reveals lasting functional changes in the regenerative output of xenografted human HSC exposed to dual inflammatory stress despite both the 10w recovery period and the unaffected HSC frequency.

### Identification of two transcriptionally and epigenetically distinct HSC compartments

To understand the molecular basis of the decreased human graft size observed with dual inflammatory challenge, immunophenotypic CD45+CD34+CD38-CD45RA-human stem/early progenitor cells were isolated from NSG xenografts at 20w and subjected to scMultiome profiling (Fig. 2A). Cell-type assignment of 27,492 single cells was guided by projection and classification via BoneMarrowMap (fig. S5, A to C, and Table S5). Notably, cell composition by transcriptional identity did not fully align with the expected downstream progenitor composition within the sorted population; most committed progenitors are typically excluded (*17*). Rather, transcriptomes corresponding to immunophenotypic CD38+ progenitors including granulocyte macrophage progenitors (GMP), megakaryocyte erythrocyte progenitors (MEP), and common lymphoid progenitors (CLP) were identified. This suggests that freshly isolated human immunophenotypic early progenitor cells in this inflammation recovery model exhibit a degree of transcriptional variability likely induced by regeneration in the xenotransplantation model (fig. S5B).

**Fig. 2.**
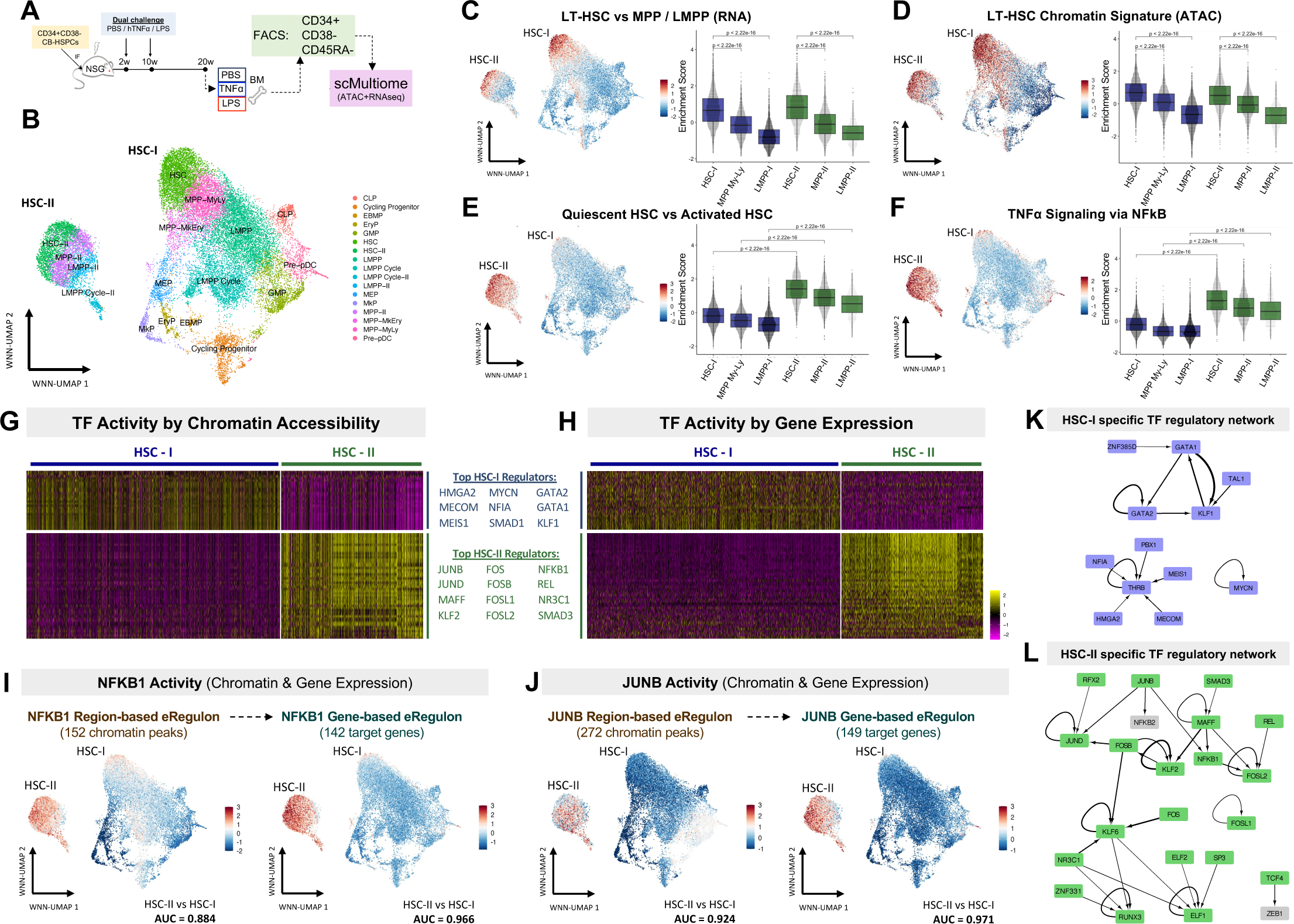
Two human HSC populations identified in an inflammation recovery model. **(A)** Schematic outlining scMultiome profiling of Prim (CD45+CD34+CD38-CD45RA-) from xenografts administered with dual inflammatory challenge. **(B)** UMAP of 27,492 xenograft HSPCs based on integrated RNA and ATAC embeddings from weighted nearest-neighbor (WNN) analysis. **(C-F)** Normalized signature enrichment scores (AUCell) of an LT-HSC-specific gene (C) or chromatin (D) signature from purified cell fractions, quiescent vs. activated HSC signature (E) and a TNFα via NF-kB signaling hallmark geneset (F) overlaid on the WNN UMAP and also depicted as boxplots for the indicated populations. Statistical comparisons are made with a Wilcoxon rank-sum test. **(G-H)** Enrichment of transcription factor (TF) activity in HSC-I or HSC-II based on target chromatin accessibility (G) and target gene expression (H) inferred using SCENIC+. TFs in rows are matched in both heatmaps. Top regulators for each HSC state are also indicated. **(I-J)** Inferred TF activity of NFKB1 (I) and JUNB (J) by target chromatin peak accessibility and target gene expression overlaid on the xenograft WNN UMAP. AUC scores for discriminatory power between HSC-I and HSC-II are also depicted for each TF activity score. **(K-L)** TF regulatory networks specific to either HSC-I (K) or HSC-II (L) from SCENIC+. TFs with significant enrichment by both target gene expression and target chromatin accessibility activity in HSC-I or HSC-II are colored in blue or green, respectively. Regulation strength between TFs is represented by edge width.

Unexpectedly, we observed two populations among cells classified as bona fide HSCs, denoted HSC-I and HSC-II (Fig. 2B, and fig. S5, D to F). Separate MPP and lympho-myeloid primed progenitor (LMPP) populations were identified contiguous with HSC-II and designated as MPP-II and LMPP-II. Both HSC-I and HSC-II express stem cell-specific marker genes relative to their downstream MPP and LMPP populations including *MECOM, AVP*, and *CRHBP* (fig. S5G) (*15, 21, 50*). Principal component analysis (PCA) of gene expression showed HSC-I and HSC-II could be discriminated from downstream progenitors using PC1 (fig. S5, H and I). Crucially, both HSC-I and HSC-II displayed enrichment of LT-HSC-specific gene expression (Fig. 2C, and fig. S5, J and K) and chromatin accessibility (Fig. 2D) signatures derived from prior bulk sequencing datasets of purified HSPC (*21*), in addition to matching HSC reference transcriptomes (fig. S5I). Thus, we confidently confirm their identity as HSCs.

Multiple computational approaches were used to compare HSC-I to HSC-II. First, weighted nearest neighbor analysis suggested that HSC-I and HSC-II represent distinct transcriptional and epigenetic states in the context of the inflammation-recovery model (Fig. 2B, and fig. S5, E and F). Parallel analyses with alternative scRNA and scATAC analysis pipelines including OCAT (fig. S6, A to C) (*51*), TooManyCells (fig. S6, D to F) (*52*), and TooManyPeaks (fig. S6, G to I) (*53*) provided further support that HSC-I and HSC-II represent distinct cellular states. Indeed, differential accessibility (DA) analysis between HSC-I and HSC-II revealed 2,112 differentially accessible regions (DARs) specific to HSC-II (fig. S7A, and Table S6), as well as global changes in TF binding site accessibility (fig. S7B) and specific enrichment for JUN/FOS motifs from the AP-1 family (fig. S7, C and D) in HSC-II. Furthermore, differential expression (DE) analysis revealed 2,178 genes upregulated in HSC-II (fig. S7, E to G, and Table S7), along with enrichment of signatures corresponding to human LT-HSC quiescence (Fig. 2E), TNFα via NFkB signaling (Fig. 2F), and TGF-ꞵ signaling (fig. S7, H and I). Importantly, HSC-I and HSC-II also separated along defined PCA and latent semantic indexing (LSI) components from dimensionality reduction of gene expression and chromatin accessibility, respectively (fig. S7, J to M). Finally, pathway analysis revealed diverse biological pathways significantly enriched in HSC-II, spanning immune signaling, proteostasis and stress responses, and regulation of cell cycle and cell motility, whereas no biological pathways were significantly enriched in differentially expressed genes (DEGs) specific to HSC-I (fig. S7N). Thus, HSC-I and HSC-II represent distinct cell states as defined by global gene expression and chromatin accessibility differences.

We next sought to understand differences between HSC-I and HSC-II using gene regulatory networks (GRNs). SCENIC+ (*54*) was utilized to integrate paired gene expression and chromatin accessibility information from individual cells, and construct “e-Regulons’’ to identify key TFs and GRNs underlying HSC-I and HSC-II states (Fig. 2, G and H, and fig. S8, A to C, and Table S8). The GRN governing HSC-I included stemness regulators *HMGA2, MECOM, PBX1*, and *MEIS1*, and regulators of megakaryocyte-erythroid fate including *GATA2, GATA1*, and *KLF1* (Fig. 2K) (*55*). GRNs governing HSC-II included inflammatory TFs *NFKB1* (Fig. 2I, and fig. S8D) and *REL* (fig. S8F), AP-1 family members *JUNB* (Fig. 2J, and fig. S8E), *JUND, FOS, FOSB, FOSL1, FOSL2* (fig. S8G), and *MAFF* (fig. S8H), among others (Fig. 2L). Collectively, these data provide strong evidence that the two human HSC populations observed within our inflammation-recovery model represent transcriptionally and epigenetically distinct states, with HSC-II distinguished primarily by enrichment of inflammatory stress programs.

### HSC-II harbor memory of prior inflammatory stress

Since inflammatory TF GRNs are active within HSC-II, it was important to interrogate molecular changes resulting from hTNFα or LPS treatments in each HSC subset separately. Therefore, we compared the transcriptional and epigenetic differences between HSC-I and HSC-II after recovery from hTNFα or LPS treatment compared to PBS-treated control xenografts. The extent of these differences became more pronounced after hTNFα or LPS treatment (Fig. 3A, and fig. S7, J to M). Furthermore, quantification with Augur (*56*) revealed greater separability of HSC-I and HSC-II by gene expression and chromatin accessibility following hTNFα or LPS challenge compared to PBS controls (Fig. 3B).

**Fig. 3.**
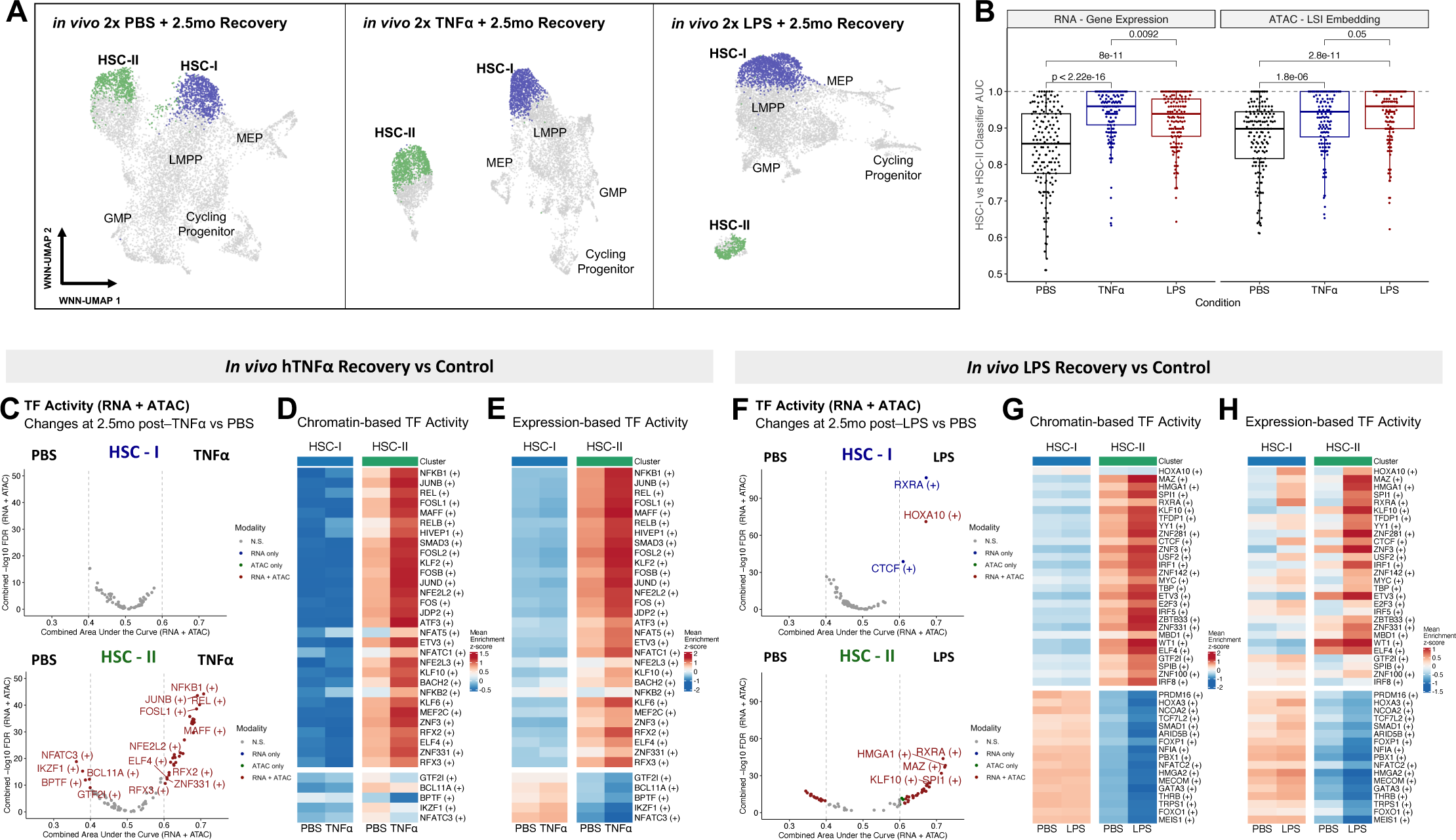
HSC-II retains memory of prior inflammatory stress. **(A)** WNN UMAPs of xenografted Prim HSPCs by dual inflammatory treatment condition, namely PBS (n = 9,756), hTNFα (n = 8,785), or LPS (n = 8,951). **(B)** Separability of HSC-I and HSC-II within each experimental condition using Augur. Each point represents the AUC of a random forest classifier trained to predict HSC-I vs HSC-II status from 150 subsamples of the data. Summary data is presented as box plots with a line at the median. Distributions were compared using a Wilcoxon rank-sum test. **(C-E)** Differential SCENIC+ TF activity between *in vivo* dual hTNFα challenge compared to PBS control, stratified by HSC subset. Colored points in the volcano plots (C) have significance at FDR < 0.05 and an AUC < 0.4 or AUC > 0.6. TFs that met these criteria by both RNA and ATAC are colored red. Mean scores for differentially enriched TFs by chromatin accessibility (D) or gene expression (E) is depicted for each condition and each HSC subset. **(F-H)** Differential SCENIC+ TF activity between *in vivo* dual LPS challenge compared to PBS control, presented as in (C-E).

To investigate the transcriptional and epigenetic changes present in either HSC-I and HSC-II as a consequence of inflammatory stress, we performed differential enrichment analysis of epigenetically and transcriptionally inferred TF activity from SCENIC+. HSC-II cells displayed higher activity of 30 TFs after recovery from hTNFα compared to PBS controls, including *NFKB1* and *JUNB*, on the basis of both target region accessibility and target gene expression (Fig. 3, C to E, and Table S9). By contrast, HSC-I cells exhibited no differences in TF activity after recovery from hTNFα treatment beyond an AUC specificity threshold of 0.6 (Fig. 3C). Differences in TF activity comparing HSCs after LPS recovery to PBS controls also showed the most pronounced differences in HSC-II, where activity of 28 TFs were enriched, including *HMGA1, SPI1, IRF1*, and *MYC* (Fig. 3, F to H, and Table S9). However, unlike hTNFα-challenged HSCs, upregulation of *HOXA10, RXRA* and *CTCF* activity was observed specifically in HSC-I following LPS recovery (Fig. 3, F to H, and Table S9); other TFs did not consistently differ in activity at the level of both RNA and chromatin in HSC-I. In summary, epigenetic and transcriptional changes in HSCs following recovery from inflammation occurred primarily within the HSC-II compartment, suggesting that HSC-II are preferentially impacted by, and retain memory of prior inflammatory stress. Given these findings, we termed HSC-II as inflammatory memory HSCs (HSC-iM) (*57–59*).

### HSC-iM share core molecular programs with memory T cells

Next, we asked if the transcriptional and epigenetic programmes in HSC-iM may be broadly shared with immune cells from two classical models of immunological memory: BCG vaccination and T cell memory. There was no enrichment of the HSC-iM gene expression signature in either murine or human HSCs “trained” by BCG vaccination (fig. S9, A to D) (*60, 61*). Similarly, gene expression signatures of murine or human HSCs following BCG vaccination failed to distinguish HSC-iM from HSC-I (fig. S9, E to F, and Table S10; AUCs of 0.597 and 0.436, respectively).

T cell memory is a well studied example where the AP-1 family TFs (*62, 63*), which are highly specific to the HSC-iM state (Fig. 4B), are also known to play a key role. We re-analyzed RNA-seq and ATAC-seq data from a study where antigen-specific CD8 memory T cells (T-mem) and CD8 effector T cells (T-eff) were purified from volunteers 4-13 years after yellow fever vaccination and deuterium ingestion, alongside naive CD8 T cells (naive-T) purified prior to vaccination (Fig. 4A) (*64*). There was high concordance in motif enrichment (R=0.77, *p*<2.2e-16) when comparing open chromatin regions specific to long-term T-mem against those specific to HSC-iM (Fig. 4B). In particular, AP-1 family TF binding sites defined by a core 7-bp motif - 5’-TGAG/CTCA-3’ - were highly enriched in accessible chromatin regions specific to both HSC-iM and T-mem (Fig. 4B). Conversely, TBR-2 (*EOMES*) and TBX21 (T-bet) motifs were specific to T-mem, as expected (*65*). In addition, open chromatin regions specific to HSC-iM were enriched in T-mem (Fig. 4, C and D) compared to T-eff (*p*=0.012) and naive-T (*p*=0.0044). Similarly, the transcriptional signature specific to HSC-iM (Fig. 4E) was enriched in T-mem relative to both T-eff (*p*=0.036) and naive-T (*p*=0.0043; Fig. 4F); enrichment was confirmed by GSEA (fig. S10A). Conversely, HSC-I signatures were not enriched in T-mem (fig. S10, B and C). A T-mem transcriptional signature comprising 257 genes upregulated in T-mem when compared to both naive-T and T-eff cells (Fig. 4G) was specifically enriched in HSC-iM compared to HSC-I (Fig. 4H, and Table S10; AUC=0.964). The specificity was more pronounced within the hTNFα-(AUC=0.987) and LPS-treated (AUC=0.965) conditions compared to PBS (AUC=0.913) (fig. S10D). A core set of 35 overlapping genes was identified by comparing the 257 gene T-mem signature and the top 200 HSC-iM marker genes; these 35 genes were capable of identifying memory subsets within both HSC and T cells (fig. S10, E and F, and Table S10). Independently derived CD4 and CD8 memory T cell signatures from a CITE-seq dataset (fig. S11, A to K) (*66*) were both also enriched in HSC-iM, moreso following recovery from hTNFα and LPS compared to PBS (fig. S11L, and Table S10). Taken together, these data indicate that there is convergence between epigenetic and transcriptional programs underlying inflammatory memory in human HSCs and functionally-defined human T cell immune memory.

**Fig. 4.**
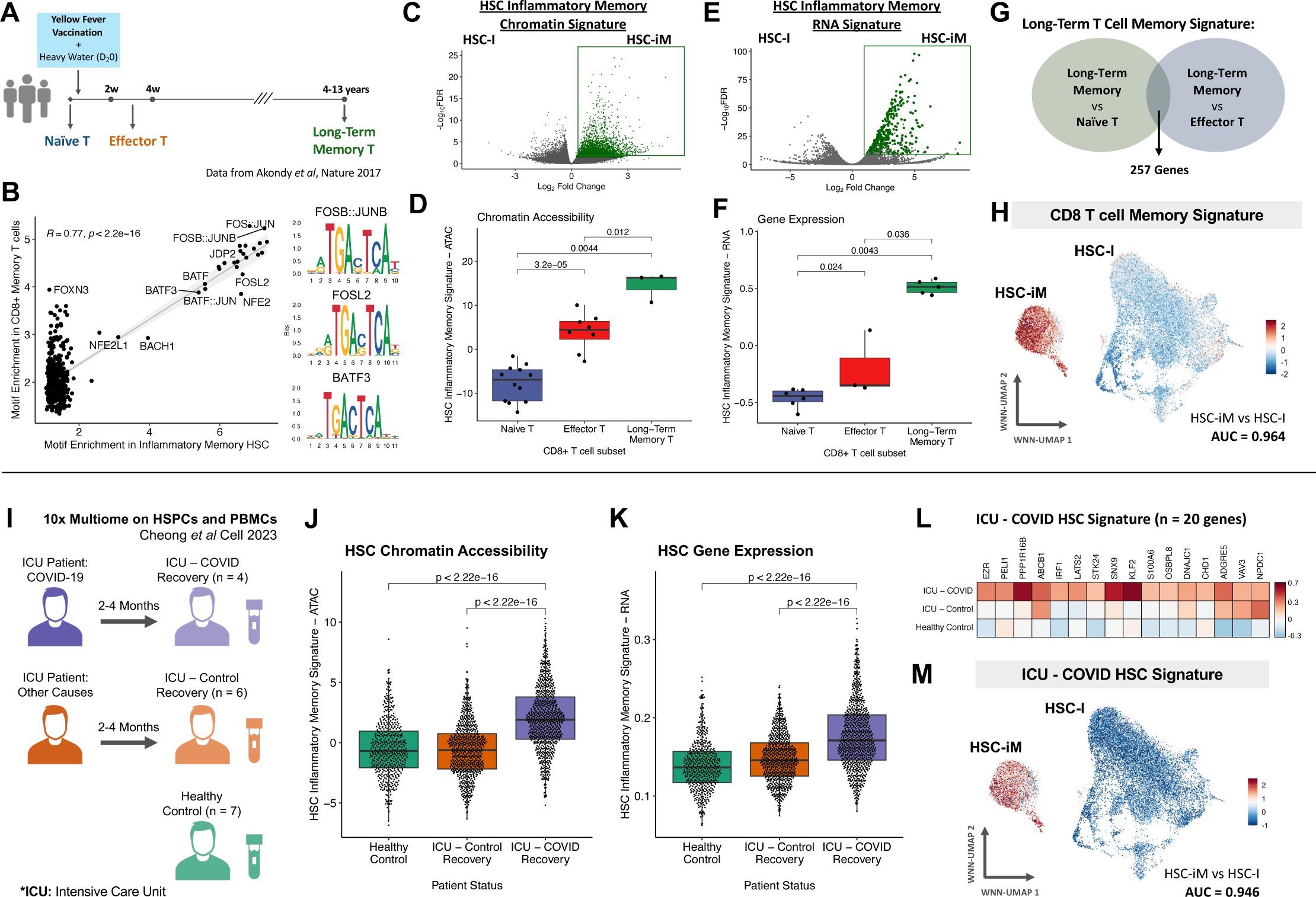
HSC inflammatory memory transcriptionally resembles human T cell memory and is induced by COVID-19 infection. **(A)** Schematic outlining purification of CD8 T cell subsets from human volunteers **(B)** Pearson correlation of TF motif enrichment among differentially accessible regions (DARs) specific to memory T cells vs HSC-iM. Three co-enriched motifs are shown as examples. **(C-D)** Enrichment of an epigenetic HSC-iM (vs HSC-I) signature comprising 3,663 DARs (C) was evaluated within human T cell subsets (D). **(E-F)** Enrichment of a gene expression-based signature (E) composed of the Top 200 differentially expressed genes (DEGs) specific to HSC-iM (vs HSC-I) was evaluated within human T cell subsets (F). **(G)** Derivation of a human CD8 long-term T cell memory signature of 257 significantly enriched (FDR < 0.01, LFC > 1) genes. **(H)** Scoring of the CD8 long-term T cell memory signature in xenograft scMultiome data. AUC score for discriminatory power between HSC-I and HSC-iM is also shown. **(I)** Experimental setup of scMultiome profiling of HSPC from donors 2-4 months following severe COVID-19 infection, as well as ICU recovery controls and healthy donors. **(J-K)** Enrichment of HSC-iM signatures within human HSCs from donors in (I), scored on the basis of chromatin accessibility (J) and gene expression (K). **(L)** Mean normalized expression of 20 DEGs comprising a post-COVID HSC signature upregulated in HSCs from ICU-COVID donors compared to healthy controls and ICU-controls at FDR < 0.05. **(M)** Scoring of the post-COVID HSC signature in xenograft scMultiome data. AUC score for discriminatory power between HSC-I and HSC-iM is also shown. Statistical comparisons, where indicated, by Wilcoxon rank-sum tests.

### HSC inflammatory memory programs are enriched in patients who recovered from severe COVID-19

We examined the relevance of our HSC-iM program in the context of patients that recovered from severe COVID-19 infections, as COVID-19 can trigger a cytokine storm and uncontrolled inflammatory responses, leading to acute respiratory distress syndrome and requiring intensive care (*45, 67*). We evaluated scMultiome-profiled HSPCs collected from patients 2-4 months following intensive care unit (ICU) admission for either severe COVID-19 (ICU-COVID; n=4) or other causes (ICU-control; n=6), alongside HSPCs from healthy donors (healthy control; n=7; Fig. 4I) (*67*). Using BoneMarrowMap, we identified 3,759 HSCs *in silico* (fig. S12A). Each clinical condition was evaluated for enrichment of the HSC-iM signature at the level of both chromatin accessibility and gene expression. The strongest enrichment was found in HSCs from ICU-COVID patients compared to HSCs from healthy controls or ICU-controls (Fig. 4, J and K, and fig. S12B). In contrast, the HSC-I signature was negatively enriched in HSCs from ICU-COVID patients compared to HSCs from healthy controls or ICU-controls (fig. S12, C and D). Moreover, a post-COVID HSC transcriptional program comprising 20 DEGs unique to ICU-COVID HSCs compared to healthy control HSCs and ICU-control HSCs (Fig. 4L) was specifically enriched in HSC-iM compared to HSC-I (Fig. 4M, and Table S10; AUC=0.946). This specificity was modestly more pronounced within the hTNFα (AUC=0.957) and LPS (AUC=0.949) recovery conditions compared to PBS control (AUC=0.915) (fig. S12E). Together, these data demonstrate that the HSC-iM signature is relevant to a real-world setting of recovery from severe inflammation in humans.

### HSC inflammatory memory accumulates in human HSC with age

Repeated inflammatory insults, including infections, occur throughout the human lifespan. As “inflammaging” - chronic, low-grade inflammation - is associated with physiological aging (*68*), we asked whether the HSC-iM signature accumulates in human HSCs with age. To this end, we analyzed 23,048 HSC transcriptomes across 4 distinct BM cohorts encompassing a total of 41 donors between 19 and 87 years of age (fig. S13, A to D). The HSC-iM signature was significantly enriched in HSCs from older-aged (OA) or middle-aged (MA) donors compared to HSCs from young adult (YA) donors by GSEA (Fig. 5A). Furthermore, DE genes unique to aged HSCs were significantly enriched in HSC-iM relative to HSC-I by GSEA (Fig. 5B, and Table S11). In contrast, an HSC-I gene signature exhibited variable patterns of enrichment between young and aged HSCs in a dataset-dependent manner (fig. S13E). Similarly, HSCs from MA and OA donors had stronger enrichment for the HSC-iM chromatin signature and lower enrichment for an HSC-I chromatin signature compared to HSCs from YA donors (fig. S13, F and G).

**Fig. 5.**
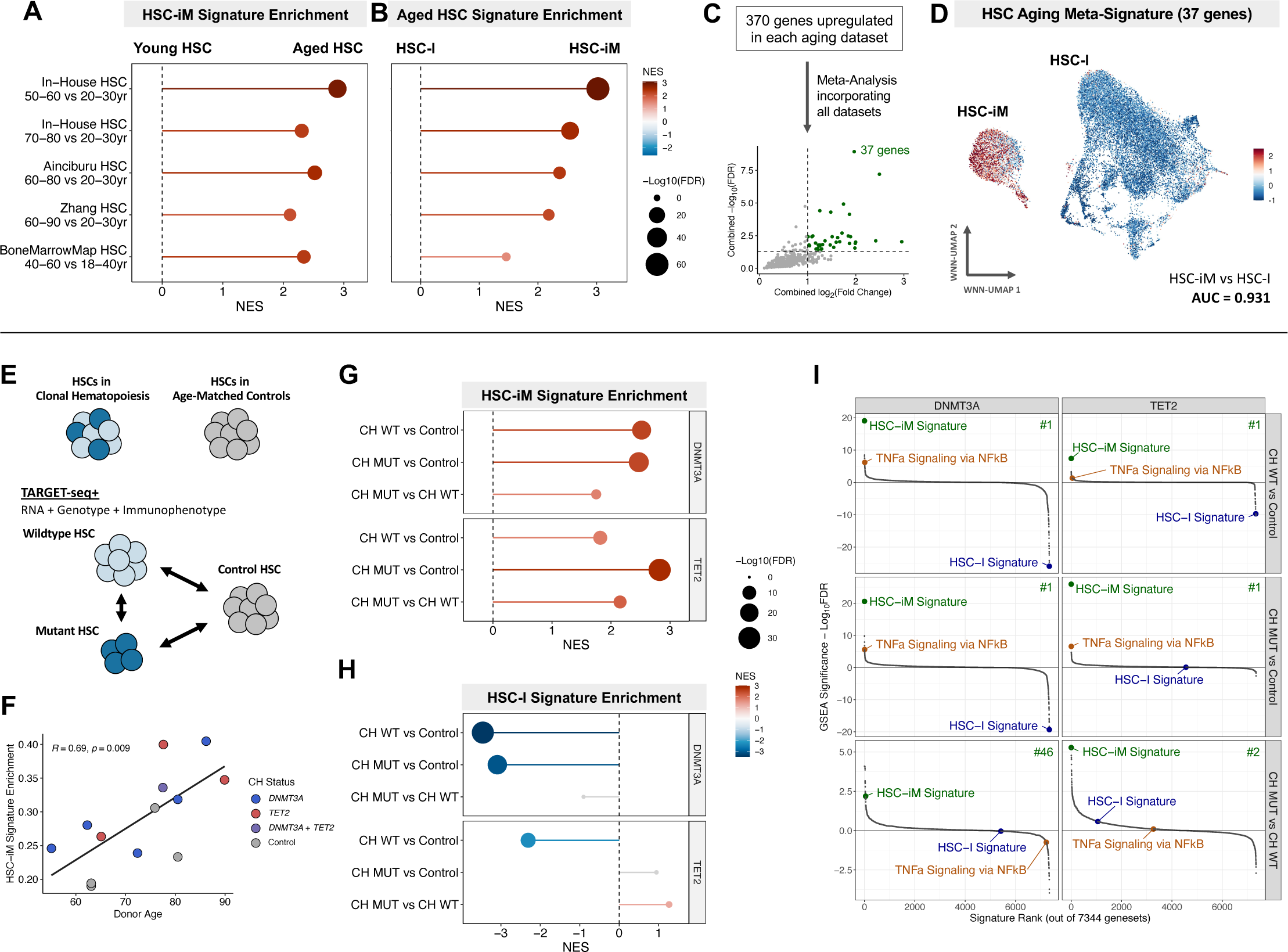
HSC inflammatory memory programs accumulate with human aging and clonal hematopoiesis. **(A)** GSEA results depicting HSC-iM gene expression signature enrichment in differential expression results between five separate comparisons of older-aged (OA; 60-90y) or middle-aged (MA; 40-60y) vs young adult (YA; 18-40y) human HSCs. **(B)** GSEA results depicting enrichment of individual Aged HSC signatures from datasets in (A) in xenograft HSC-iM vs HSC-I **(C-D)** Human HSC aging datasets from (A-B) were integrated to derive a 37-gene Aged HSC-specific meta-signature (C) which was scored in the xenograft scMultiome data (D). AUC score for discriminatory power between HSC-I and HSC-iM is also depicted. **(E)** Schematic outlining TARGET-Seq+ profiling of HSPCs from nine CH donors and four non-CH controls. **(F)** Pearson correlation between donor age and HSC-iM signature enrichment within 4,651 transcriptional HSCs from CH and non-CH donors. **(G-H)** GSEA results depicting HSC-iM (G) and HSC-I (H) signature enrichment in differential expression results between Control, CH^WT^, and CH^MUT^ HSC/MPPs, after adjusting for multiple covariates. Positive and negative enrichment indicate upregulation and downregulation in CH, respectively. Results are faceted by CH mutation type. **(I)** Benchmarking of GSEA results from HSC-iM and HSC-I signatures against signatures from MSigDB, totaling 7,344 genesets. The enrichment rank of the HSC-iM signature is depicted in the top right corner of each comparison. Results are faceted by CH mutation type.

To minimize effects of donor- and dataset-driven variability, we defined a human HSC aging meta-signature of 37 genes consistently upregulated across all comparisons and datasets, including *NR4A1* which has been linked to inflammation-resistance in CH (*69*) (See Methods; Fig. 5C, and fig. S13H, and Table S11). This signature was highly enriched in HSC-iM compared to HSC-I at the single cell level (AUC = 0.931; Fig. 5D and fig. S13I). Enrichment was more pronounced within the hTNFα-(AUC=0.971) and LPS-treated (AUC=0.932) conditions compared to PBS control (AUC=0.848; fig. S13J). These findings demonstrate that the transcriptional profile of the HSC-iM state accumulates in HSCs during human aging.

### HSC inflammatory memory is linked to clonal hematopoiesis

The association of the HSC-iM signature with human HSC aging prompted us to interrogate inflammatory memory in CH, a condition that is associated with both aging (*70, 71*) and elevated serum levels of inflammatory cytokines (*29*). We took advantage of TARGET-seq+ data from our study of nine *DNMT3A* or *TET2*-mutated CH donors and four age-matched controls with no detectable CH mutations (Fig. 5E) (*72*). TARGET-seq+ combines simultaneous sc-profiling of transcriptome, genotype, and cell surface immunophenotype, enabling comparisons of mutant (CH^MUT^) and wild-type (CH^WT^) HSCs obtained from individuals with CH to HSCs from age-matched controls without CH (control non-CH; fig. S14A). Of note, CH^WT^ and control non-CH are operational terms in relation to the variants tested, and we cannot rule out the possibility of undetected mosaic chromosomal alterations or mutations in unknown CH driver genes.

Among 4,651 transcriptionally-defined HSCs (fig. S14, B to C), enrichment of the HSC-iM signature was correlated with age across the thirteen donors (R=0.69, *p*=0.0085, Fig. 5F). As age and CH are closely tied, we investigated whether the presence of expanded CH clones is also linked to HSC-iM. HSCs from CH donors were compared to HSCs from control non-CH donors after adjusting for donor ID, age, sex, and cell sorting batch as covariates (Fig. 5E). Unexpectedly, both CH^WT^ and CH^MUT^ HSCs from the *DNMT3A-* and *TET2-*mutated CH donors were significantly enriched for the HSC-iM signature compared to control non-CH HSCs (Fig. 5G). Conversely, the HSC-I signature showed decreased enrichment in both CH^WT^ and CH^MUT^ HSCs (Fig. 5H). These data point to a non-cell-autonomous role for HSC inflammatory memory in the context of CH.

In order to benchmark the relative strength of these associations, we extended our GSEA by incorporating additional biological genesets from MSigDB (totaling 7,344 genesets) and ranked the enrichment results based on statistical significance. Strikingly, enrichment for the HSC-iM signature surpassed all other genesets, including TNFα via NFkB signaling, for all comparisons: CH^WT^ or CH^MUT^ HSCs from donors with *DNMT3A-* or *TET2*-mutated CH against control non-CH HSCs (Fig. 5I). Regulons constructed from SCENIC analyses that were upregulated in CH^WT^ HSC compared to control non-CH HSCs included several TFs associated with inflammatory memory: *NFKB1, RELB*, and AP-1 family members, *JUNB* and *FOSL1* (fig. S14D). Moreover, a signature of 219 genes transcriptionally upregulated in CH^WT^ HSCs was highly enriched in HSC-iM compared to HSC-I at the single cell level (AUC = 0.945; fig. S14, E to G, and Table S12), and this enrichment was more pronounced within the hTNFα (AUC=0.968) and LPS (AUC=0.923) inflammation-recovery conditions compared to PBS control (AUC=0.908; fig. S14H). Within individual CH donors, the HSC-iM signature was enriched in CH^MUT^ compared to CH^WT^ HSCs (Fig. 5I), although the strength of these associations was more modest; the HSC-iM signature was ranked #46 and #2 out of 7,344 genesets evaluated in HSC from *DNMT3A-* and *TET2-*mutated donors, respectively. These data indicate that presence of *DNMT3A* and *TET2* CH mutations potentiates the transcriptional inflammatory memory response in HSCs. In summary, we found that the HSC-iM state is enriched in HSCs from individuals with CH regardless of whether the mutation was present, implying that molecular changes stemming from HSC inflammatory memory underlie both non-cell-autonomous and autonomous changes in CH^WT^ and CH^MUT^ HSC, respectively.

### Identification of xenograft-derived HSC-iM signatures in human CH and aging samples

Given the strength of enrichment patterns for HSC-iM in the context of CH, our CH data set presented a unique opportunity to address whether HSC-I and HSC-iM (as identified in Fig. 2 within the xenograft setting) can be identified in an unbiased manner from human adults. In our contemporaneous CH study (*72*) where functionally validated immunophenotypic markers (*73*) were used to isolate HSPCs, we had performed unsupervised feature selection and clustering using the Self-Assembling Manifolds (SAM) tool on primitive HSPC transcriptomes, which revealed three clusters of human HSC within the cohort, termed HSC1, HSC2, and HSC3 (Fig. 6A, and fig. S15, A and B). The ratio of HSC2 to HSC1 cells was higher among CH donors compared to control non-CH donors (*p*=0.046; Fig. 6, B and C). Notably, all of the donors in our CH study had osteoarthritis, which is associated with low-grade inflammation (*74*). Similar to HSC-iM, HSC2 was enriched for an LT-HSC quiescence vs activation signature and TNFα via NFkB signaling (fig. S15, C and D). HSC1 was highly enriched for the xenograft-derived HSC-I signature (vs HSC2, AUC=0.944), while HSC2 was highly enriched for the HSC-iM signature (vs HSC1, AUC=0.963; Fig. 6D, and Table S12). Conversely, a gene expression signature representing HSC1 from the CH dataset (fig. S15E) mapped with high specificity to the HSC-I population from the xenograft dataset (vs HSC-iM, AUC=0.964), and a signature representing HSC2 (fig. S15F) was highly specific to HSC-iM (vs HSC-I, AUC=0.992; Fig. 6E, and Table S11). Notably, a core signature comprised of 67 genes with shared expression in HSC-iM and HSC2 effectively separated HSC1 and HSC2 clusters within individual BM donors (fig. S15, G and H), and distinguished HSC-iM from HSC-I within the xenograft model (fig. S15I, and Table S11).

**Fig. 6.**
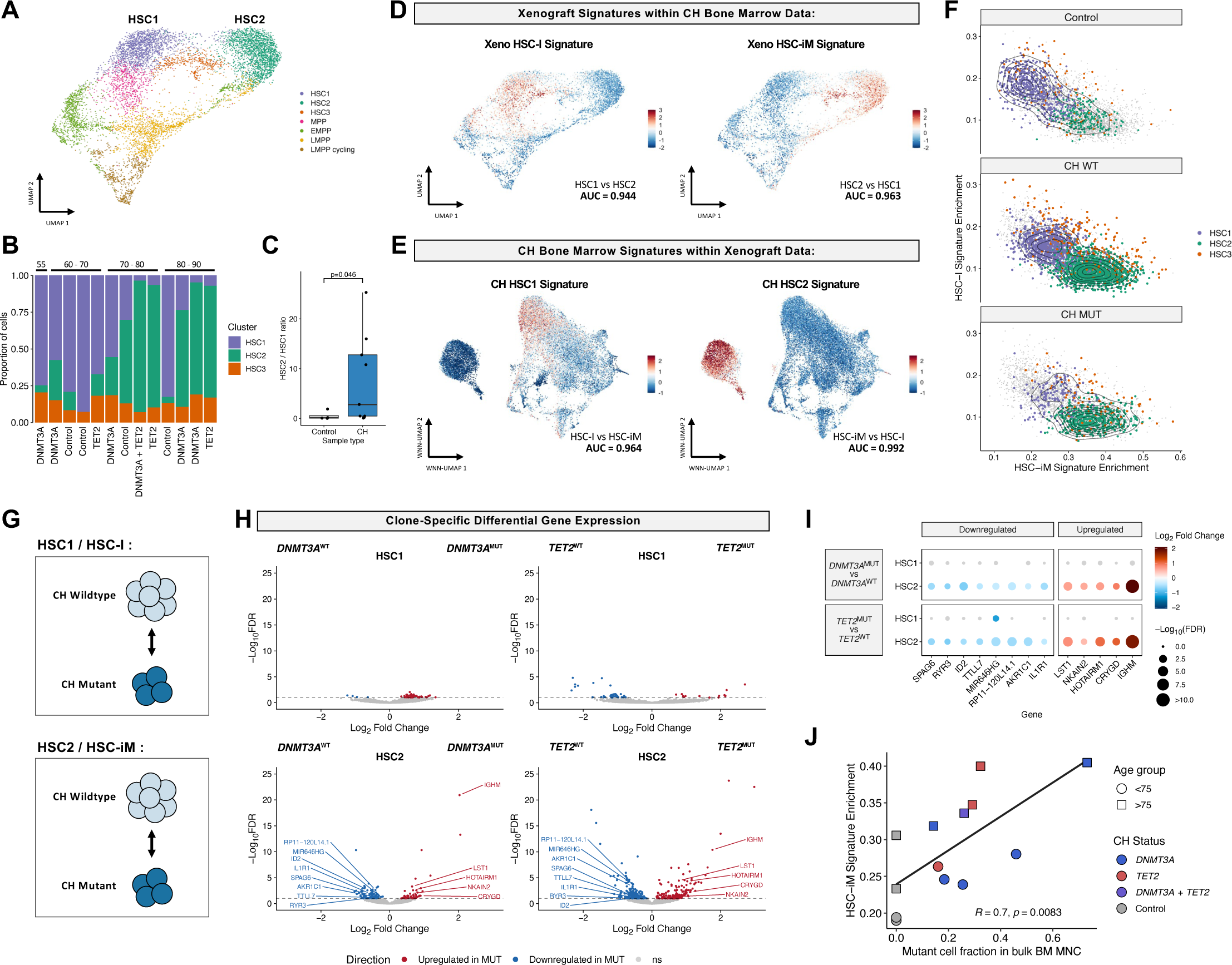
Clone-specific gene expression changes in CH are observed predominantly in HSC-iM. **(A)** UMAP of 8,059 bone marrow HSPCs from 9 donors with CH and 4 age-matched samples without CH, after unsupervised feature selection and clustering using the Self-Assembling Manifolds (SAM) algorithm. **(B)** Proportion of cells in the HSC1, HSC2, and HSC3 clusters by donor. **(C)** Ratio of HSC2 to HSC1 cells in donors with or without CH. Statistical comparison was made using a *t-*test. **(D)** Scoring of the xenograft HSC-I and HSC-iM signatures in CH BM HSPC data. AUC score for discriminatory power between HSC1 and HSC2 is also depicted for these signatures. **(E)** Scoring of the CH BM HSC1 and HSC2 cluster signatures in xenograft data. AUC score for discriminatory power between HSC-I and HSC-iM is also depicted. **(F)** Enrichment of HSC-I and HSC-iM signatures in cells belonging to each of the three HSC clusters from Control and CH donors separated by genotype. **(G)** Schematic showing the strategy for differential expression analysis between CH^MUT^ and CH^WT^ cells in either the HSC1 or HSC2 clusters. **(H)** Differentially expressed genes between CH^MUT^ and CH^WT^ cells within the HSC1 and HSC2 clusters. Genes are colored based on significance at FDR < 0.05. **(I)** Differential TF activity inferred by SCENIC between CH^MUT^ and CH^WT^ cells in either the HSC1 or HSC2 cluster. Y-axis portrays the significance level of differential enrichment by a linear mixed model accounting for donor identity. TFs are colored based on significance at FDR < 0.05. **(J)** Pearson correlation between HSC-iM signature enrichment within transcriptional HSCs and mutant cell fraction in bulk mononuclear cells (MNCs) from CH and non-CH donors.

We next analyzed HSC-I and HSC-iM signature enrichment in cells belonging to each of the three HSC clusters from non-CH and CH donors separated by CH^WT^ or CH^MUT^ status. Control non-CH, CH^WT^, and CH^MUT^ HSCs span a transcriptional gradient of increasing HSC-iM signature enrichment (Fig. 6F). HSC3 exhibited intermediate enrichment of both HSC-I and HSC-iM signatures, potentially representing a transitional population between these key HSC states (Fig. 6F). Notably, there was an overall shift in signature strength within CH^MUT^ HSC towards the HSC-iM state compared to other conditions, underscoring the relevance of inflammatory memory in CH (Fig. 6F). Together, our data provide strong evidence that HSC1 and HSC2, identified from TARGET-seq+ profiles of BM HSPCs from CH donors and non-CH controls, are equivalent to the HSC-I and HSC-iM subsets identified by 10x scMultiome profiling of xenografted CB HSPCs.

To determine whether CH mutations impacted HSC1 and HSC2 differently, we compared gene expression between CH^MUT^ and CH^WT^ HSC from both *DNMT3A-* or *TET2*-mutated donors within each HSC subset (Fig. 6G). Gene expression changes between CH^MUT^ and CH^WT^ cells occurred predominantly within HSC2 for both *DNMT3A-* and *TET2-*mutated donors (Fig. 6H, and fig. S15J). Furthermore, SCENIC showed that dysregulated TF activity between CH^MUT^ and CH^WT^ was exclusive to HSC2 and not observed in HSC1 (fig. S15, K and L) (*72*). Consistent with findings that *TET2* and *DNMT3A* mutations exert distinct epigenetic effects (*75, 76*), we only identified 13 conserved mutant gene-specific DEGs in HSCs from *TET2* and *DNMT3A* donors (Fig. 6I, and Table S13). Notably, downregulation of inflammatory genes *IL1R1* and *ID2* within CH^MUT^ HSC2/HSC-iM was found, in line with observed downregulation of TNFα via NFkB signaling within this population (*72*). These results are similar to those from the *in vivo* inflammation-recovery model wherein molecular changes following inflammatory treatment occurred predominantly within the HSC-iM population (Fig. 3, C to H). Notably, we found that the enrichment of the HSC-iM signature was correlated with CH clone size in bulk bone marrow sequencing data (R=0.70, *p*=0.0083, Fig. 6J) suggesting that the overall effect of the CH mutation on HSC-iM underlies its clonal advantage. Collectively, these data show that transcriptional dysregulation enacted by CH mutations occurs predominantly within the HSC-iM state.

## Discussion

Here, we report the discovery of a new HSC subset, termed HSC-iM, that retains transcriptional and epigenetic memory of prior inflammation. The HSC-iM molecular program is highly concordant with signatures underlying human T cell memory, pointing to convergence between these disparate cell types in their response to repeated inflammatory stimuli. The recently reported inflammatory memory in epidermal stem cells provides support for the concept of conserved inflammatory memory programs amongst stem cells (*57–59*). What is remarkable is that although HSC-iM were identified using human CB cells in a xenograft inflammation-recovery model, the molecular programs underlying HSC-iM are conserved in three human physiological settings: in HSCs from patients following severe COVID-19 illness; in HSCs as humans age; and in HSCs from people with CH. We propose that within the heterogeneous human HSC pool, the HSC-iM subset has a heritable function as a sensor of inflammatory insults. Thus, HSC-iM provides a crucial cellular connection between infection and inflammatory history, human aging, and clonal disorders of the hematopoietic system.

In mouse models, inflammation activates stem cells in the acute setting and severely disrupts stem cell fitness in the chronic setting (*14, 36*). Age-related inflammatory change is a well-recognized risk factor for many diseases, including cancer, cardiovascular disease, chronic liver disease, autoimmune disease, and diabetes (*70, 71, 77–79*). Aging is also the greatest risk factor for clonal expansion of tissue stem cells (*80, 81*), with CH being the most studied exemplar (*70, 71, 79*). CH has a prevalence of approximately 1-in-5 in people over 60 years of age (*70, 71, 79*) and represents a pre-leukemic phase of myeloid blood malignancies (*27*). Independent of age, CH results in a 12-fold increased risk for blood cancers, a 2-fold increased risk for cardiovascular disease, and other diseases associated with human aging (*70, 71, 78, 79*). Although some CH mutations are known to upregulate inflammatory cytokines (*29, 78, 82*), it was unclear prior to our study how inflammation, occurring prior to acquisition of CH mutations, drives deleterious molecular changes within HSCs. Moreover, the identity of the HSC subset that is impacted by CH mutations as well as the mechanisms that underlie the selection of CH mutant clones were also not clear.

By establishing an inflammation-recovery xenotransplantation model able to recapitulate HSC states found in aging and CH, we have identified clues towards answering these important questions. Critically, the HSC-iM subset identified from our xenograft model is remarkably similar, at a molecular level, to an HSC subset found in adult BM from our separate CH study (*72*). We found that *DNMT3A* and *TET2* mutations have both cell-autonomous and non-cell-autonomous effects on HSCs from CH donors. First, gene expression changes in CH mutant cells occur predominantly within HSC-iM/HSC2. Second, the finding that HSC-iM enrichment occurs in both CH^MUT^ and CH^WT^ HSCs suggests that individuals with CH either had greater exposure to, or were more impacted by inflammaging. Third, the data presented here and in our companion study (*72*) showed that while the overall HSC-iM signature was highest in CH^MUT^ HSC, some components of this signature related to inflammatory pathways, especially TNFα via NFkB signaling, were decreased in CH^MUT^ compared to CH^WT^ HSCs.

Collectively, these data suggest a mechanism that might explain why a CH mutation provides a clonal advantage to mutation-bearing HSCs. We propose that the HSC-iM subset emerges as a ‘cost’ of adaptation to inflammation, where the modification to HSC fitness has both beneficial and deleterious consequences (*83, 84*). HSCs are wired to prevent damage from being propagated; in some settings like low-dose radiation or proteostatic stress, damaged HSCs are efficiently culled (*85, 86*). We hypothesize that repeated inflammatory stress drives the affected HSCs into dormancy as a protective adaptation, providing beneficial effects by safeguarding the integrity of the stem cell pool over a lifetime (*83, 87*). Support for this concept comes from a distinguishing feature of HSC-iM, where their transcriptional and epigenetic state is concomitantly enriched for both an inflamation response signature and a LT-HSC signature enriched for quiescence and depleted for activation (Fig. 2, E and F) (*23*). We extend our hypothesis to propose that CH mutations attenuate at least some of the deleterious effects imposed by repeated age-related inflammatory challenges within the HSC-iM subset, for example, by reduced NFkB signaling after TNFα exposure. This results in clonal advantage for the CH mutation-bearing HSC-iM. This hypothesis is driven by our finding that HSC-iM signature enrichment correlates with CH clone size across all our samples. Hence, HSC-iM represents a stem cell reservoir that shields HSCs from damage caused by inflammatory events that occur during aging. But this adaptation at the organismal level could potentially impact human health due to the negative effects resulting from the promotion of HSC dormancy, resulting in the loss of an active HSC pool, as well as enhanced differentiation of abnormal myeloid cells with altered pro-inflammatory function and increased risk of disease (*27, 32, 88*). We further speculate that these pro-inflammatory and mutated myeloid progeny of mutant CH HSCs participate in a positive feedback loop for selection of CH mutant HSCs by providing an enhanced inflammatory milieu. In summary, an HSC subset that retains memory of prior inflammatory stress through heritable molecular alterations provides a cellular mechanism that begins to explain why health outcomes due to aging and aging-associated human diseases are heterogeneous.

## Acknowledgments

We thank the obstetrics unit of Trillium Health, William Osler and Credit Valley Hospitals for CB; K.B.K, Q.L., A.M., J.E. and N.M. for technical help; The UHN-Sickkids Flow Cytometry facility for cell sorting and PMGC for 10x scMultiome library prep and sequencing; all Dick lab members for manuscript feedback.

## Funding

University of Toronto MD/PhD studentship award (AGXZ)

Medical Research Council and Leukaemia UK Clinical Research Training Fellowship MR/R002258/1 (NAJ)

Princess Margaret Cancer Foundation (SZX, JED)

Ontario Institute for Cancer Research through funding provided by the Government of Ontario (JED)Canadian Institutes for Health Research RN380110-409786 (JED)

International Development Research Centre Ottawa Canada (JED)

Canadian Cancer Society 703212 (JED)

Terry Fox New Frontiers Program project grant 1106 (JED)

University of Toronto’s Medicine by Design initiative with funding from the Canada First Research Excellence Fund (JED)

The Ontario Ministry of Health (JED)

Canada Research Chair (JED)Medical Research Council Molecular Haematology Unit Programme Grant MC_UU_00029/8 funding 574 (PV)

Blood Cancer UK Programme Continuity Grant 13008 (PV)

NIHR Senior Fellowship (PV)

The Oxford Biomedical Research Centre Haematology Theme (PV)

## Author contributions

Conceptualization: SZX

Methodology: SZX, AGXZ, MSN, NAJ, ST

Investigation: SZX, AGXZ, MSN, INXL, LJ, JA, DP, JM, HK, MB, EW

Formal analysis: AGXZ, MSN, NAJ, SS, AM, JGC, LZ, HSL,SZX

Visualization: AGXZ, MSN, NAJ, SS, LZ, HSL

Resources: AGAN, EFF, LDS

Supervision: BW, GWS, SZJ, PV, JED, SZX

Funding acquisition: PV, JED, SZX

Writing - Original Draft: AGXZ, MSN, JED, SZX

Writing - Review & Editing: all authors.

