## Supplemental Materials for "Identification of a human hematopoietic stem cell subset that retains memory of inflammatory stress"

Biological Materials and Methods

Computational Methods

Supplementary References

Supplemental Figures S1 to S15

### Biological Materials and Methods

Human cord blood (CB) samples
Human CB samples were obtained with informed consent from Trillium Health, Credit Valley and William Osler Hospitals (Toronto, ON) according to procedures approved by the University Health Network (UHN) Research Ethics Board. Mononuclear cells (MNC) from pools of male and female CB (~1-8 donors) were obtained by centrifugation over a density barrier of Lymphoprep (Multicell, Canada). After red cell lysis with ammonium chloride (STEMCELL, Vancouver, BC) MNCs were enriched for hematopoietic stem and progenitor cells (HSPCs) by positive selection with the CD34 Microbead kit (Miltenyi, Germany) per manufacturer's instructions, or by lineage depletion as described (*1*). Resulting HSPC-enriched CB cells were cryopreserved in 50% PBS, 40% fetal bovine serum (FBS) and 10% DMSO and stored at -80°C short-term or -150°C long-term.

CB LT-HSC scRNA-seq
Human immunophenotypic long-term hematopoietic stem cells (LT-HSCs) were purified from pooled CB samples based on a Lin-CD34+CD38-CD45RA-CD90+CD49f+ immunophenotype (see FACS section below). CB LT-HSCs were purified and submitted to the Hospital for Sick Children Genomics Core for scRNA-seq profiling on the BD Rhapsody platform.

Mice
Animal experiments were done in accordance with institutional guidelines approved by the UHN Animal Care Committee. The following mouse strains (The Jackson Laboratory, Bar Harbor, ME) were used in this study: NOD.Cg-$Prkdc^{scid}Il2rg^{tm1Wjl}/SzJ$ (NSG; strain 005557), NOD.Cg-$Prkdc^{scid}Il2rg^{tm1Wjl}$Tg(CMV-IL3,CSF2,KITLG)1Eav/MloySzJ (NSG-SGM3; strain 013062), and NOD.Cg-$Prkdc^{scid}Il2rg^{tm1Wjl}Kit^{em1Mvw}$/SzJ (NSGW41). All *in vivo* experiments were done with 8- to 12-week-old female or male mice. NSG and NSG-SGM3 were conditioned with 225cGy of gamma radiation; NSGW41 females were not irradiated. All mice were housed at the animal facility (ARC) at Princess Margaret Cancer Centre in a room designated only for immunocompromised mice with individually ventilated racks equipped with complete sterile microisolator caging (IVC), on corn-cob bedding and supplied with environmental enrichment in the form of a red house/tube and a cotton nestlet. Cages were changed at least once a week under a biological safety cabinet. Health status was monitored using a combination of environmental monitoring and evaluation of soiled bedding from sentinel mice.

Fluorescence-activated cell sorting (FACS) and flow cytometry analyses
HSPC-enriched human CB cells were thawed via slow dropwise addition of X-VIVO 10 media (LONZA, Switzerland) with 50% FBS and 100µg/mL DNaseI (Roche, Switzerland). Cells were centrifuged at 400g for 10min, then resuspended in PBS+5% FBS. The populations sorted for individual experiments are indicated; populations are designated based on the full stem and progenitor hierarchy as described previously (*2*, *3*). Cells were resuspended at $<10 \times 10^6$ cells/mL and stained in one or two subsequent rounds for 15min at room temperature each. See Table S14 for antibodies. Cells were washed following staining and resuspended in PBS+5%FBS and filtered through 40µm nylon mesh for sorting or analyses. To isolate xenografted human cells, bone marrow from 3-5 mice per experimental group was thawed, pooled, mouse depleted using a MACS-based kit (Miltenyi) and stained for cell sorting. Cells were sorted on BD FACSAria

Fusion, BD FACSAria III or BD Symphony S6 cell sorters, or analyzed on BD FACSCelesta or BD Symphony A1 coupled to a high-throughput system (HTS).

Xenotransplantation

Following cell sorting, 8,000-10,000 CD34+CD38- cells were transplanted by intrafemoral injection into the right femur of appropriately conditioned age- and sex-matched mice in a volume of 30μL PBS. To induce acute inflammatory stress, mice were injected intraperitoneally with 5μg hTNFα or 40μg lipopolysaccharide (LPS) at the indicated time points. Control groups were injected with the vehicle control PBS. Mice were euthanized at the indicated time points by cervical dislocation and the injected femur along with other hindlimb long bones (non-injected femur and both tibias) collected into Iscove's modified Dulbecco's medium (IMDM)+5%FBS. Spleens were also collected in the same medium. Injected and non-injected bones were flushed separately in IMDM+5%FBS using a 23g needle for some experiments. Alternatively, bones from most mice euthanized at a 20w post-transplant time point were crushed in PBS+5%FBS using a mortar and pestle and the resulting cell suspension passed through a 40-70μm nylon mesh. Spleens were macerated using a syringe plunger, then passed through a 40μm nylon mesh. Typically, 5-10% of cells were analyzed for human chimerism and lineage markers using flow cytometry; antibodies are listed in Table S14. Remaining cells were cryopreserved. For serial transplantation, xenografted cells were sorted for human CD45 (see Table S14) and transplanted into NSG-SGM3 mice at the indicated cell doses. A mouse was considered engrafted if human chimerism was >0.1% and LT-HSC frequency by limiting-dilution assay (LDA) was estimated using the ELDA software (http://bioinf.wehi.edu.au/software/elda/) (*4*).

T cell culture and CapTCRseq

Approximately 300,000 CD45+CD34-CD19-CD33- were isolated by FACS as described above from cryopreserved xenograft bone marrow to enrich T cells and deplete B, myeloid and progenitor cells. Cells were seeded in X-VIVO 20 media supplemented with 5% human AB serum and 30U/mL hIL-2 and stimulated with anti-CD3/CD28 Dynabeads (0.5 beads per cell) (*5*). Live cells were counted by trypan blue exclusion on a hemocytometer and maintained in culture for 8 days before harvesting. Genomic DNA was extracted with a QIAamp DNA blood mini kit (Qiagen), quantified using a NanoDrop ONE instrument, and submitted for CapTCRseq (*6*) at the Princess Margaret Genomics Centre. Shannon's diversity index was calculated for the reads obtained for the three TCR genes as described previously (*7*).

scMultiome sorting and library preparation

For scMultiome profiling, cryopreserved bone marrow from xenografted mice was processed for FACS as described above and sorted for CD19-CD34+CD38-CD45RA- live cells. CB and human bone marrow was similarly thawed and stained for FACS and sorted for CD19-CD34+CD38- and CD19-CD34+CD38+ populations. The two populations were pooled back together at known ratios for further processing. Sample processing, including nuclei isolation and library construction, was performed by the Princess Margaret Genome Centre for downstream 10x Genomics scMultiome RNA + ATAC sequencing using their standard or low-input protocol, as appropriate.

### Computational Methods

CB LT-HSC scRNA-seq QC and preprocessing

Raw sequencing data was aligned using the BD SevenBridges pipeline, with default parameters. CB LT-HSCs were demultiplexed from other samples in the same run based on sample specific barcodes. Cells passing the following criteria were used for downstream analysis: unique feature counts > 500, total number of molecules detected within a cell > 1000, percent mitochondrial genes < 5%. Raw counts were normalized using scran (v1.20.1) (*8*), followed by variable gene selection, scaling, PCA, and neighbor graph construction with 10 PCA components and k-nearest neighbors equal to 10. The neighbor graph was used to construct a force-directed graph for visualization using 1.8.2 Scanpy (*9*).

Consensus non-negative matrix factorization

Consensus non-negative matrix factorization (cNMF) (*10*) was run with 100 iterations on the single cells from components 2-15 to pull out transcriptional gene programs of variation in an unsupervised manner. Raw counts were fed into the –counts flag, and SCTransform V2 (*11*) corrected counts specifically from the data slot of the SCT assay were fed into the –tpm flag of the cNMF command line function with 10 workers. After running cNMF across various components, the optimal number of components was selected by assessing the silhouette score (stability) and Frobenius reconstruction error (error), as implemented in cNMF. The transcriptomic signatures were characterized using an overrepresentation analysis with the function, 'fora' from the package fgsea (v1.18.0) (*12*) on the top 100 genes from each signature with HSC genesets and additional pathways from MSigDb.

scMultiome (RNA and ATAC) preprocessing

Chromium Single-Cell Multiome ATAC + Gene Expression sequencing data (scMultiome) was pre-processed with CellRanger-ARC (v2.0.0) and aligned to GRCh38. The RNA feature count matrix for each sample was corrected for ambient RNA contamination using SoupX (v1.6.2) (*13*) with default parameters. SoupX-corrected RNA counts were processed using Seurat (v4.3.0) (*14*) and ATAC peak counts were processed using Signac (v1.6.0) (*15*). RNA and ATAC quality control (QC) filtering utilized the following metrics: percentage of mitochondrial genes (pct_mito), nucleosome banding pattern (nucleosome_signal), ATAC transcriptional start site enrichment (TSS_enrichment), number of unique RNA transcripts detected (nFeature_SoupX), and number of fragments in peaks (nCount_ATAC). Thresholds for QC filtering were adapted to each dataset based on the distribution of QC metrics. For the CB Xenograft scMultiome samples, filtering thresholds were set at: nFeature_SoupX > 1000, pct_mito < 18%, nCount_ATAC > 1000, nucleosome_signal < 2, TSS_enrichment > 1. For the primary CB scMultiome samples, filtering thresholds were set at: nFeature_SoupX > 300, pct_mito < 8%, nCount_ATAC > 1000, nucleosome_signal < 2, TSS_enrichment > 1. For the Aging scMultiome samples, filtering thresholds for RNA-seq were set unique to each sample: BM_24M (nFeature_SoupX > 300, pct_mito < 10%), BM_26F (nFeature_SoupX > 200, pct_mito < 15%), BM_57M_58F (nFeature_SoupX > 100, pct_mito < 18%), BM_70F (nFeature_SoupX > 200, pct_mito < 12%), BM_77F (nFeature_SoupX > 200, pct_mito < 14%). For the Aging scMultiome samples, ATAC-seq filtering thresholds were uniform for each sample: nCount_ATAC > 1000, nucleosome_signal < 2, TSS_enrichment > 1.

Furthermore, pooled scMultiome samples with multiple donors (CB pools A and B, BM Middle-Aged donors, CB Xenograft) were separated using SoupOrCell (*16*) through genotype-based clustering of sequencing reads from scRNA-seq profiles. The optimal number of genotypic clusters was selected using an elbow plot of the total log probability, or manually selected when the number of donors was known.

Donor Sex Classification

Pools of CB donors were engrafted into NSG mice from which CB Xenograft samples were derived. Further, some libraries of primary CB and BM samples consisted of a pool of one male sample and one female sample. To account for variation from donor sex in CB xenograft samples and to demultiplex pools of primary CB and BM samples, we developed an approach to cluster single cells by donor sex.

High-confidence sex classifications were first assigned by ATAC and RNA. For ATAC: among cells with an ATAC read depth of >6000, those with one or more ChrY reads were classified as Male, and those without were classified as Female. For RNA: cells were scored for male-specific (on ChrY, excluding the pseudo-autosomal region) genes and female-specific (XIST and TSIX) genes using AddModuleScore from Seurat (v4.3.0). These scores were min-max normalized between 0 and 1, and adaptive thresholding was applied using a function adapted from the AUCell (v1.14.0) (*17*) package. Cells surpassing this enrichment threshold for the male-specific geneset with zero female-specific gene expression were classified as Male, and those surpassing this enrichment threshold for the female-specific geneset enrichment with zero male-specific gene expression were classified as Female. Cells confidently classified as Male or Female by both RNA and ATAC were used, and differential expression was performed between these Male and Female cells to identify donor-specific genes within each dataset. Dimensionality reduction and unsupervised clustering was subsequently performed based on these donor-specific genes to cluster cells into male and female donors, and to identify clusters of doublets expressing both male and female genes. This approach was validated on artificially mixed snRNA-seq data from male and female HSPC from CB and BM samples, and applied to multiplexed scMultiome samples from CB Xenograft, primary CB, and primary BM.

Doublet Identification

Within multiplexed samples, doublet cells with mixed donor genotypes were identified by SoupOrCell (*16*). Simultaneously, snRNA-seq based doublet identification was performed using scDblFinder (v1.6.0) (*18*) with default parameters. We confirmed that transcriptomes with co-expression of male-specific and female-specific genes were captured as doublets by SoupOrCell as well as scDblFinder. All identified doublets were removed from downstream analysis.

CB Xenograft Processing and Classification

After initial quality control, single-cell transcriptomes from the CB Xenograft experiment were normalized by scran (v1.20.1), followed by variable feature selection, scaling, and PCA reduction. Batch correction with harmony (v0.1.1) (*19*) was performed based on inferred donor sex. Initial dimensionality reduction and clustering revealed two outlier clusters each representing < 1% of all cells, appearing to be contaminating T cells and pro-B cells. These clusters were excluded, and this data was re-processed, using the top 20 harmony-corrected principal components, which were used to construct a neighborhood graph using the top 30 neighbors for each cell. UMAP reduction was performed with min.dist=0.2 and spread=1.

For chromatin analyses, peaks were called using MACS2 (*20*) from fragment files corresponding to high quality cells after QC filtering. Peaks were called independently from HSPCs in each treatment condition (PBS + recovery, TNF + recovery, LPS + recovery), following the Signac pipeline. Peaks within non-standard chromosomes, genomic blacklisted regions, and those spanning less than 20bp or more than 10,000bp were removed from downstream analyses. From the resulting peak matrix, dimensionality reduction with latent semantic indexing (LSI) was performed following the Signac pipeline and LSI components were batch corrected by donor sex using harmony. Harmony-corrected LSI components 2 to 30 were used to construct a neighborhood graph using the top 30 neighbors for each cell. UMAP reduction was performed with min.dist=0.2 and spread=1.

For integrative analysis, weighted nearest neighbors (WNN) integration (*21*) was performed using harmony-corrected RNA PCA components 1:20 and harmony-corrected ATAC LSI components 2:30, considering the top 30 neighbors for each cell. UMAP reduction was performed with a min.dist=0.2 along the WNN neighborhood graph.

<u>CB Xenograft cell state annotations</u>
For cell state annotations, transcriptomes were projected onto BoneMarrowMap (https://github.com/andygxzeng/BoneMarrowMap) and filtered at a mapping error threshold of 2 MADs above the median. These cell state assignments were used to guide clustering from the WNN graph: leiden clustering was performed at varying resolutions from 0.1 up to 10. Specific leiden clusters from resolutions of 1, 5, or 10 were selected based on high concordance with cell state assignments from BoneMarrowMap, and a combination of these clusters was used for cell state annotation. For a subset of cells (~1%) that were not captured by these clusters, annotation was performed based on the most frequent cell state annotation of their nearest neighbors.

<u>Integrated UMAP embeddings by CB treatment condition</u>
For embeddings generated within each condition, WNN integration was run using the top 30 neighbors from harmony-corrected RNA PCA components 1:20 and harmony-corrected ATAC LSI components 2:20. UMAP reduction was performed from neighborhood graphs for RNA only, ATAC only, and integrated WNN with min.dist = 0.2 and spread = 1.2. This was repeated for each individual condition: PBS + Recovery, TNF + Recovery, LPS + Recovery.

<u>OCAT embedding by CB treatment condition</u>
The conditions “PBS”, “TNF” and “LPS” were each treated as a separate batch. The three batches of scRNA-seq data commonly share 36,601 genes, each with 10,097 cells, 8,069 cells, and 8,100 cells, respectively. No genes or cells were filtered in this analysis. OCAT (*22*) was used to integrate these three batches of scRNA-seq datasets. OCAT pre-processed the raw gene expression matrices through log-transformation and l2-normalization and performed dimension reduction on the original data to a d=100 subspace. OCAT then selected 100 “ghost” cells, centers of small cell neighborhood in the reduced subspace, and connected each individual cells to the global “ghost” cell set (m=100*3) through a bipartite graph. OCAT further made these

edge weights sparse by allowing at most 30% of them to be non-zero. These sparse edge weights are treated as the OCAT representation of each cell and were used for downstream visualization with UMAP.

TooManyCells embedding by CB treatment condition

For analysis with TooManyCells (*23*), single-cell transcriptomes were filtered to remove cells with less than 250 counts and only include genes present in at least 1 cell to discard low-quality reads. The filtered samples were normalized with term frequency-inverse document frequency (tf-idf) and clustered with the TooManyCells divisive hierarchical clustering using matrix-free spectral clustering. The resulting cluster trees were pruned to a minimum of 30 cells for each leaf node to reduce the overall size of the tree and focus visualization on larger sub-populations of cells. Once pruned, the trees were labeled with cell-type annotations, highlighting the clustered differences between HSC and HSC-II populations, including additional annotation for MPP-MyLy (associated with the HSC population) and MPP-II (associated with the HSC-II population).

TooManyPeaks embedding by CB treatment condition

For analysis with TooManyPeaks (*24*), chromatin peak matrices derived from scATAC-seq were filtered to remove cells with less than 1000 peaks. The filtered samples were processed with latent semantic analysis for dimensionality reduction with 50 components. The peaks for scATAC-seq were not normalized. The filtered samples were clustered with TooManyPeaks, the sister method to TooManyCells. The resulting clustered trees were pruned to a minimum of 200 cells, as scATAC-seq data usually produces an exceptionally large number of clusters (leaf nodes). Once pruned, the trees were labeled in the same manner as the scRNA-seq data analyzed with TooManyCells.

Marker gene derivation within CB Xenograft scMultiome data

To find reliable markers for each cell state, a composite score was derived from four distinct differential expression statistics. At the single cell level, the wilcoxauc function from presto (v1.0.0) (*25*) was applied to obtain the following statistics: 1) log2 Fold Change (logFC) from single cells in group vs single cells from all other groups, 2) AUC metric for distinguishing between cells in group vs cells from all other groups. Next, pseudo-bulk profiles were created by combining cells based on CellType, Donor Sex, and Treatment condition (excluded pseudobulks with <5 cells) and DESeq2 (*26*) was used to compare pseudobulks from each cell state against those from all other cell states through an likelihood ratio test (LRT). Based on this pseudo-bulk analysis, the following statistics were also incorporated: 3) test statistic from DESeq2, and 4) logFC from DESeq2. The geometric mean from all four statistics was used to prioritize marker genes for each cell state, and subsequently called “MarkerScore”.

This metric was also used to identify top markers between HSC-I and HSC-II, although the pseudo-bulk DESeq2 analysis was performed directly contrasting HSC-I and HSC-II, and accounting for treatment (PBS, TNF, LPS) as a covariate. The top 200 genes for each population was used as a signature for scoring in external datasets.

Differential chromatin accessibility within CB Xenograft scMultiome data

For differential chromatin accessibility, pseudo-bulk profiles were created by combining scATAC-seq cells based on CellType, Donor Sex, and Treatment condition and DESeq2 (*26*) was used to compare pseudobulks from each cell state against those from all other cell states through an likelihood ratio test (LRT), while adjusting for treatment (PBS, TNF, LPS) as a covariate. Significantly differentially enriched peaks were used as signatures for scoring enrichment in external datasets. Among differentially enriched peak sets, TF motif enrichment was determined through the "FindMotifs" function in Signac (v1.6.0)

Gene set enrichment analysis

Gene set enrichment analysis (GSEA) was performed using the function fgseaMultilevel from fgsea (v1.18.0) (*12*), with the following parameters: nPermSimple=1000000, eps=0, minSize=15, maxSize=500. The test statistic from DESeq2 (*26*) was used as the rank statistic for each gene. For complete biological pathway analysis, GSEA was performed using the April 2023 version of "Human_GOBP_AllPathways_no_GO_iea" from Gary Bader's lab in Toronto (https://download.baderlab.org/EM_Genesets/current_release/Human/symbol/). EnrichmentMap (*27*) was used for visualization of biological pathway GSEA results, only retaining signatures and pathways enriched in a given cell type at FDR $< 0.01$.

Augur

Augur (*28*) was used to determine separability of single cells between HSC-I and HSC-II pertaining to gene expression and chromatin accessibility profiles, applied independently to each treatment condition (PBS, TNF, LPS). For gene expression, augur was applied on the normalized expression of 2000 highly variable genes, with default parameters. Whereas, for chromatin accessibility, augur was applied to 50 LSI components, with default parameters. For ATAC, these reduced dimension components were used rather than raw chromatin peak counts due to high sparsity and number of features for the latter. The distribution of augur classifier performance for each cross-validation split was used to represent the separability between HSC-I and HSC-II within each treatment condition.

RNA and ATAC signature scoring

Geneset scoring of previously published gene expression signatures was performed using AUCell (v1.14.0) (*17*). Enrichment of chromatin regions in scATAC-seq data was evaluated through chomVAR (v1.14.0) (*29*). When necessary, hg19 coordinates for chromatin signatures was converted to hg38 using package rtracklayer (v1.52.1) (*30*) with the UCSC hg19tohg38 chain file. For both AUCell (RNA) and chromVAR (ATAC) based enrichment scores, standardization was performed across cells prior to plotting for ease of visualization.

SCENIC+ e-Regulon Inference

To infer transcription factor activity from RNA and ATAC profiles by SCENIC+ (*31*) within the CB Xenograft scMultiome data, the 27,492 single cells from the xenograft scMultiome were first aggregated to the metacell level using the divide and conquer algorithm from the Metacell2 (*32*) package. The highly variable genes used to construct the uniform manifold and projection were used as input into the algorithm guiding feature gene selection, thus optimizing clustering for metacell partitioning. The approach generated 1,476 metacells with a target metacell size of

75,000 UMI. Next, pyCistopic was used to identify cell states and cis-regulatory topics from the ATAC single cells in an unsupervised manner. The ideal number of topics were evaluated as per the previously reported guidelines (*31*). Consequently, pyCistopic optimized at 18 topics. Motif enrichment analysis was subsequently carried out by leveraging pyCistarget, which utilizes precomputed databases comprising motif scores and rankings for genomic regions, and a motif-to-transcription factor annotation database from the Aerts lab resources (https://resources.aertslab.org/cistarget/databases/homo_sapiens/hg38/screen/mc_v10_clust/region_based/). This analysis was conducted on the topics from pyCistopic and differentially accessible regions between each defined cell type.

Following pyCistopic and pyCistarget, a SCENIC+ object was created for downstream steps to create enhancer-driven gene regulatory networks using the ATAC and RNA. First, regions and genes that were not present in more than 0.5% of cells were filtered out. Next, cistromes were generated using the function merge_cistromes, which overlaps targets assigned to a TF from the motif enrichment dictionaries with the regions in the object. Then, enhancer-to-gene relationships were inferred by first defining the search space around each gene of 150kb upstream/downstream, specifically on the metacells, to limit RAM usage while connecting to biomart host 98. After, enhancer-to-gene models were generated using gradient boosting machines with the function calculate_regions_to_genes_relationships. Next, TF-to-gene relationships were inferred using pySCENIC on the RNA metacells as per previously published guidelines (*33*) with a candidate list of TFs (*34*) and loaded into the SCENIC+ object from a saved adjacencies matrix using the function load_TF2G_adj_from_file. Finally, enhancer-driven gene regulatory networks were generated using a GSEA recovery approach from SCENIC+ using the function build_grn. The recovered eRegulons were filtered using the function apply_std_filtering_to_eRegulons, allowing an analysis with high confidence transcription factors.

Given that a gene expression-based eRegulon and a chromatin accessibility-based eRegulon was reported for each TF, we performed quality control by Pearson correlation and excluded TFs wherein correlation between gene expression-based eRegulon activity and chromatin accessibility-based eRegulon activity was below $r = 0.25$. This led to 133 TFs wherein eRegulon activity passed the correlation threshold and exhibited concordance between RNA and ATAC.

Differential signature enrichment

Differential enrichment of gene or chromatin signature scores was performed by constructing linear mixed models for each signature using the cell class as an independent variable and accounting for donor sex as a random effect within the data using the function “dream” from package variancePartition (v1.22.0) (*35*). The raw and FDR-corrected p values from this analysis were used to represent significance. Simultaneously, the area under the receiver operator curve (auc) metric from function “wilcoxauc” of package presto (v1.0.0) was used to represent the ability of a signature enrichment score to accurately discern between contrasting cell classes.

Applied to SCENIC eRegulons wherein two enrichment scores are used for each TF, representing activity by gene expression as well as chromatin accessibility, these two results were integrated to identify top marker TFs for a given cell type or condition. Here, the mean auc metric as well as the mean -log10(FDR) value were used together to identify top marker TFs across RNA and ATAC modalities.

SCENIC+ TF network construction

Among TFs with significant eRegulon activity enrichment in a specific cell state or condition at FDR < 0.05 by both RNA and ATAC modalities, gene regulatory networks (GRNs) were constructed. Briefly, strength of TF regulation for a target gene was represented through two metrics: 1) importance of the RNA-based TF-to-gene association, and 2) importance of the ATAC-based chromatin region-to-gene association among nearby chromatin regions containing the TF motif. The geometric mean of these two metrics was taken to represent the overall strength of regulation of target gene expression by a TF. Finally, this score was subject to adaptive thresholding in line from the AUCell package as described above, and TF-to-TF regulation surpassing this adaptive threshold was retained to construct a TF regulatory network from each condition.

Analyses of external bulk RNAseq datasets

For differential expression analysis in bulk RNAseq datasets, DESeq2 (v1.32.0) (*26*) was applied to raw count data. For sample-level scoring of geneset enrichment, GSVA (v1.40.1) (*36*) scoring was performed on normalized gene expression data with the “kcdf” parameter set to “gaussian”. Normalized data from the original studies was used when available, otherwise raw count data was subject to *vst* normalization through DESeq2. For murine bulk RNAseq data, murine genes were converted to human orthologs using babelgene (v22.3) and only conversions supported by a minimum of 5 databases were retained. These converted gene names were used for downstream processing as described.

For differential accessibility in bulk ATAC-seq datasets, peak count matrices were obtained from the original studies and differentially accessible peaks were identified through DESeq2. and scoring of chromatin signatures was performed through chromVAR. When comparing gene signature and chromatin signature enrichment between groups of samples, Wilcoxon rank-sum tests were performed.

BCG vaccination-associated transcriptional signature

To derive a signature of murine LSK+CD150+ HSCs exposed to BCG vaccine (BCG-iv) (*37*), differential expression analysis was performed through DESeq2 (*26*) comparing BCG-iv versus Control conditions. This resulted in 490 genes up-regulated in BCG-exposed murine HSCs at FDR < 0.05 and log2FC > 1, constituting a murine HSC BCG-iv signature.

To derive a signature of human CD34+CD38-CD45RA- HSC/MPPs exposed to BCG vaccine (*38*), differential expression analysis was performed through DESeq2 comparing Day 90 post-BCG vaccination versus Day 0 pre-vaccination conditions. This resulted in 206 genes upregulated in HSC/MPPs at 90 days following BCG vaccination at FDR < 0.05 and log2FC > 1, constituting a human HSC/MPP BCG vaccination signature.

‘Akondy’ CD8 Memory T cell Signature

To derive a signature of functionally defined CD8 memory T cells (*39*), differential expression was performed between Memory T (T-mem) cells and Naive T (naive-T) cells, and between T-mem and Effector T (T-eff) cells. From 937 genes enriched in T-mem versus naive-T at log2FC > 1 and FDR < 0.01 and 1543 genes enriched in T-mem versus T-eff cells at log2FC > 1 and FDR < 0.01, we identified 257 overlapping genes significantly enriched in T-mem cells compared to both naive-T and T-eff cell subsets. These 257 genes represented the CD8 T cell memory signature.

Memory T cell signature from CITEseq data

CITEseq data of human T cell subsets was obtained (*21*). Cell type annotations and UMAP coordinates were also extracted from the original publication. Differential expression was performed on pseudo-bulk profiles with DESeq2 (v1.32.0) adjusting for donor as a covariate. To derive CD8 and CD4 T cell memory signatures, DE genes unique to both Central Memory T and Effector Memory T cell subsets compared to Naive T at log2FC > 1 and FDR < 0.05 were retained, and the top 200 enriched genes were used as the signature.

Post-COVID HSC/MPP analyses and signature derivation

scMultiome of HSPCs in Healthy Control, ICU-Control, and ICU-COVID donors was profiled previously (*40*). Cells annotated as HSC/MPPs from this study were projected onto BoneMarrowMap and transcriptional HSCs were purified *in silico* by reference map projection. Differential expression analysis with DESeq2 (v1.32.0) was applied to pseudo-bulk profiles to compare HSCs from ICU-COVID donors against HSCs from ICU-Control and healthy control donors, and 20 genes enriched in ICU-COVID at FDR < 0.05 were retained as an ICU-COVID recovery molecular signature.

Differential expression between HSC across age

Profiling of primary CD34+ BM cells by scMultiome as described above yielded 43,762 cells with high quality RNA and ATAC profiles spanning 6 donors: 2 young adult (YA) donors (20-30yr), 2 middle aged (MA) donors (50-60yr), and 2 older aged (OA) donors (70-80yr). In addition to this data, scRNA-seq data was compiled from three additional cohorts comprised of bone marrow samples of varying ages: 1) 10x v2 and v3 scRNA-seq from CD34+ BM profiled in Ainciburu *et al* (*41*), spanning 5 YA donors (20-30yr) and 3 OA donors (60-80yr); QC filtering as per Jakobsen *et al* (*42*) yielded a total of 71,805 cells. 2) STRT-seq from CD34+ fractions BM profiled in Zhang *et al* (*43*) spanning 3 YA donors (20-30yr) and 2 OA donors (60-90yr); QC from the original study yielded 3,023 cells. 3) 10x v2 scRNA-seq from mixed CD34+ and bulk BM profiled in four studies (*44–48*) included within BoneMarrowMap spanning 13 YA donors (18-40yr) and 9 MA donors (40-60yr). QC filtering (filtered at nGenes > 500, pct.mito < 8%, and scrublet-based doublet filtering) yielded 194,905 cells.

Single-cell transcriptomes from each cohort were classified by BoneMarrowMap projection and transcriptional HSCs were purified *in silico* for downstream analysis. Collectively, this constituted 23,048 transcriptional HSC spanning 23 YA donors, 11 MA donors, and 7 OA donors Pseudo-bulk profiles were constructed from HSCs from each donor sample, and differential expression was performed between HSCs from older adults and HSCs from younger adults using DESeq2.

Aged HSC Meta-Signature derivation

To derive a meta-signature of aged HSCs spanning all datasets, donor-specific HSC pseudo-bulk profiles from each dataset were pooled together. Differential expression was performed between HSCs from 18 middle/older aged adults and HSCs from 23 young adults, adjusting for the originating dataset as a covariate. Prior to interpretation of differential expression results, 370 genes that were upregulated in aged HSCs at logFC > 0 within each of the four aging datasets were retained. Among these 370 universally up-regulated genes, 37 were significantly up-regulated in aged HSCs by pooled differential expression at logFC > 1 and FDR < 0.05. These 37 genes represent a meta-signature that is consistently upregulated across human HSC aging.

TARGET-seq+ clustering and CH bone marrow reference map cell type mapping

TARGET-seq+ profiling, scRNA-seq pre-processing, and cell type annotation of human BM HSPCs from clonal hematopoiesis (CH) and control donors is outlined in the original study (*42*). Gene identifiers were converted to Ensembl v93 and HSCs were annotated by projecting the dataset onto the bone marrow reference map as described above. The HSC/MPP, LMPP, LMPP cycling, and EMPP clusters from our original study 1 were further subclustered using the self-assembling manifolds (SAM) algorithm, using default settings with Harmony-adjusted PCs as input and using the sample identifier as the batch (*49*). The resulting SAM-weighted PCA was then used as input to generate a UMAP and for Louvain clustering, which identified 7 clusters.

For calculating HSC2:HSC1 cell ratio, the number of cells in each cluster was calculated on a per-sample basis. Only cells sorted as part of the total Lin–CD34+ gate were included, so as to avoid bias introduced by FACS depletion of CD38– cells.

Differential gene expression analysis of TARGET-seq+ data from CH bone marrow

Differential expression testing was performed with a linear mixed model to account for sample covariance using the dream pipeline from the variancePartition package (*35*), which is based on limma-voom (*50*). Testing was performed on log normalized counts, using the scran normalization size factors. Genes were filtered to include only those expressed in at least 10% of cells in either group. A linear mixed model was fitted to each gene using ‘dream’ and differential expression testing was performed using ‘variancePartition::eBayes’. For comparisons between CH and non-CH controls, the sample type was used as the test variable, and the sample identifier, age, sex, and FACS sort batch effects included as covariates. For comparisons between genotypes within CH samples, the clone was used as the test variable, and the sample identifier and batch effect included as mixed effect covariates. Samples were excluded from the comparison if they had less than 5 cells in either genotype.

SCENIC analysis of TARGET-seq+ data from CH bone marrow

To infer transcription factor (TF) regulon activity, regulon analysis was performed using pySCENIC (*17*), which was run as per previously published workflow guidelines (*33*). To identify candidate TF-regulons, filtered and pre-processed raw counts were used as the input alongside a list of human TFs (*34*). Candidate regulons were pruned using the annotations of TF motifs ‘motifs-v10nr_clust-nr.hgnc-m0.001-o0.0.tbl’, and CisTarget was applied using the ‘mc_v10_clust’ databases of known human TF motifs annotated at: a) 500 bp upstream and 100 bp downstream of the transcription start site (TSS); and b) 10 kilobases centered around the TSS.

No drop-out masking was applied. Enrichment of refined TF regulons was quantified using AUCell, with default parameters. Tests for differential regulon activity were performed using a linear mixed model, as described above.

Differential signature enrichment analysis of TARGET-seq+ data from CH bone marrow

Differences in gene expression signature or TF regulon scores between conditions were tested by a linear mixed model. For comparisons between CH and non-CH controls, the sample type was used as the fixed effect, and the sample identifier, age, sex, and FACS sort batch as mixed effect covariates. For comparisons between genotypes within CH samples, the clone was used as the fixed effect, and the sample identifier as a mixed effect covariate. P-values were obtained by a likelihood ratio test of the full model with the fixed effect against the model without the fixed effect.

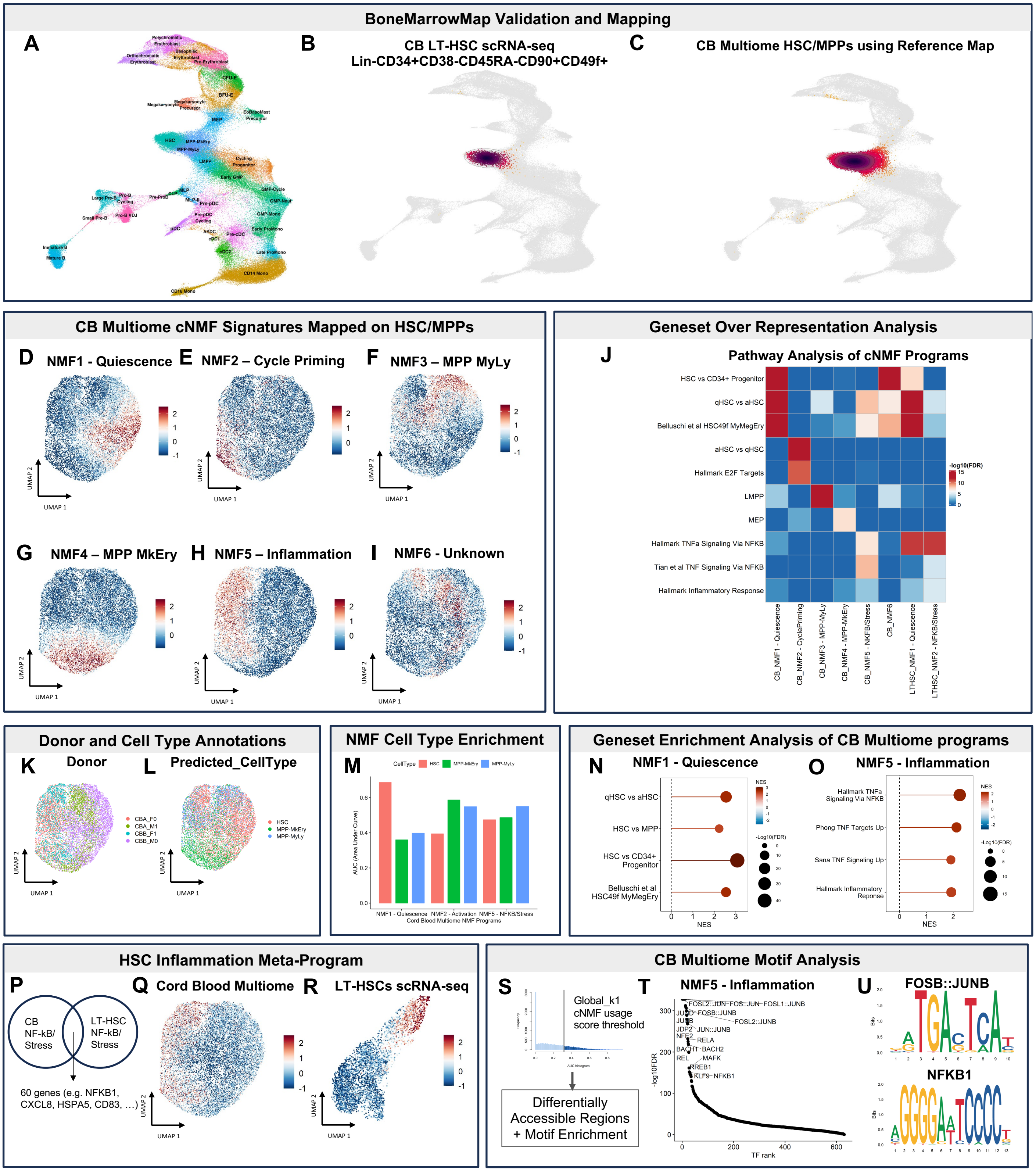


**Fig. S1. Validation of inflammatory programs in CB-HSCs with an independent scMultiome dataset.**

**(A)** Integrated UMAP embedding of the BoneMarrowMap reference atlas (B-C) Density plot of cells mapped onto BoneMarrowMap. **(B)** Single-cell RNA-seq (scRNA-seq) of Long-Term Hematopoietic Stem Cells (LT-HSCs) sorted with Lin-CD34+CD38-CD45RA-CD90+CD49f+. **(C)** Cord Blood scMultiome (CB) HSC/MPPs defined using the BoneMarrowMap **(D-I)** Enrichment of cNMF programs on a UMAP embedding of CB HSC/MPPs. **(D)** NMF1: Quiescence. **(E)** NMF2: Cycle Priming. **(F)** NMF3: MPP-MyLy. **(G)** NMF4: MPP-MkEry. **(H)** NMF5: Inflammation. **(I)** NMF6: Unknown. **(J)** Key pathways significant in cNMF programs defined on the CB scMultiome HSC/MPPs and LT- HSCs scRNA-seq. **(K-L)** Annotations mapped onto UMAP embedding of CB HSC/MPPs. **(K)** Donor. **(L)** Cell Type Annotations from reference mapping using BoneMarrowMap. **(M)** Area under the curve of cNMF programs defined on CB HSC/MPPs in each BoneMarrowMap cell type. **(N-O)** Geneset Enrichment Analysis of CB cNMF programs. **(N)** NMF1: Quiescence. **(O)** NMF5: Inflammation. **(P-R)** AUCell Enrichment of Inflammation Meta-Program. **(P)** Venn diagram of top 200 gene overlap from each respective signature. **(Q)** CB scMultiome. **(R)** LT-HSCs scRNA-seq. **(S)** Schematic outlining TF motif enrichment analysis among differentially accessible regions specific to cells that pass a threshold for the CB scMultiome NMF5: Inflammation program. **(T)** Top motifs in cells enriched with the NMF5: Inflammation program in the CB scMultiome. **(U)** Two candidate motifs from the AP-1 and NFKB family.

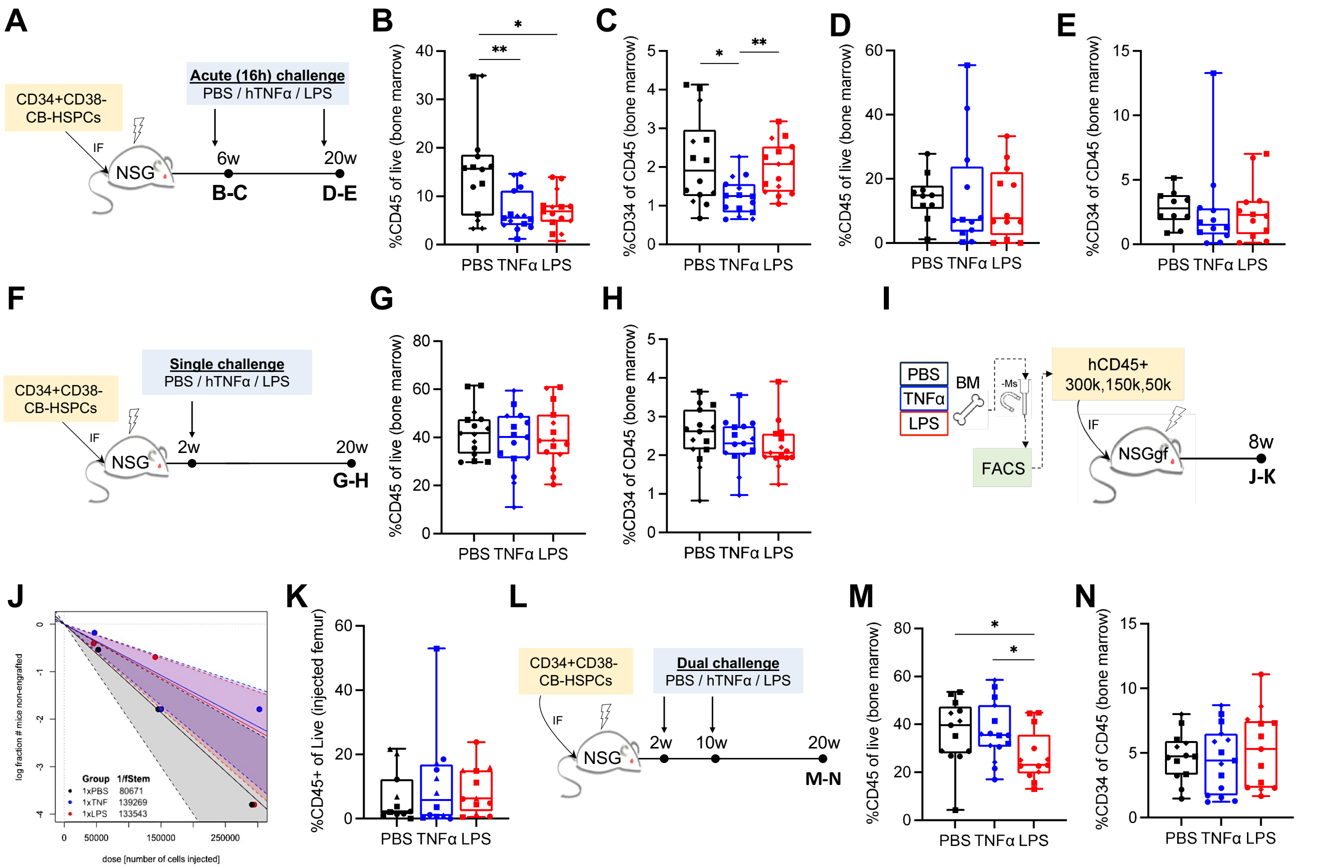

**Fig. S2. Inflammatory response and recovery of xenografted human HSCs in non-injected bone marrow.**

**(A)** Experimental schematic for acute inflammatory challenge of xenografts with hTNFα or LPS at two time points. **(B-C)** Effect of acute inflammatory challenge of xenografts 6 weeks post-transplant on human cells (B) and progenitors (C) in non-injected hind-limb bone marrow 16 hours after administration. **(D-E)** Effect of acute inflammatory challenge of xenografts 20 weeks post-transplant on human cells (D) and progenitors (E) in non-injected hind-limb bone marrow 16 hours after administration. **(F)** Schematic for xenograft model of recovery from single (2 weeks) acute inflammatory challenge. **(G-H)** Human engraftment (G) and progenitor composition (H) in non-injected hind-limb bone marrow 18 weeks after a single inflammatory challenge. **(I)** Experimental schematic for secondary transplantation of single-challenged xenografts with limiting dilution. **(J)** Stem cell frequency estimates calculated for secondary transplants in (I). No significant differences were found by chi-squared tests. **(K)** Proportion of engrafted human cells in cohorts from (I) transplanted with 300,000 hCD45+ cells. **(L)** Schematic for xenograft model of recovery from dual (2 weeks and 10 weeks) acute inflammatory challenge. **(M-N)** Human engraftment (M) and progenitor composition (N) in non-injected hind-limb bone marrow 10 weeks after dual inflammatory challenge. All engraftment data is presented as box plots with a line at the median, boxes showing quartiles and error bars showing range. Individual points show data for each animal used in the study (n=10-15), and different symbols reflect different CB pools (n=2-3). Data were compared using pairwise Mann-Whitney tests; *** p<0.001, ** p<0.01, * p<0.05, and p<0.1 is shown numerically, while p>0.1 is not shown.

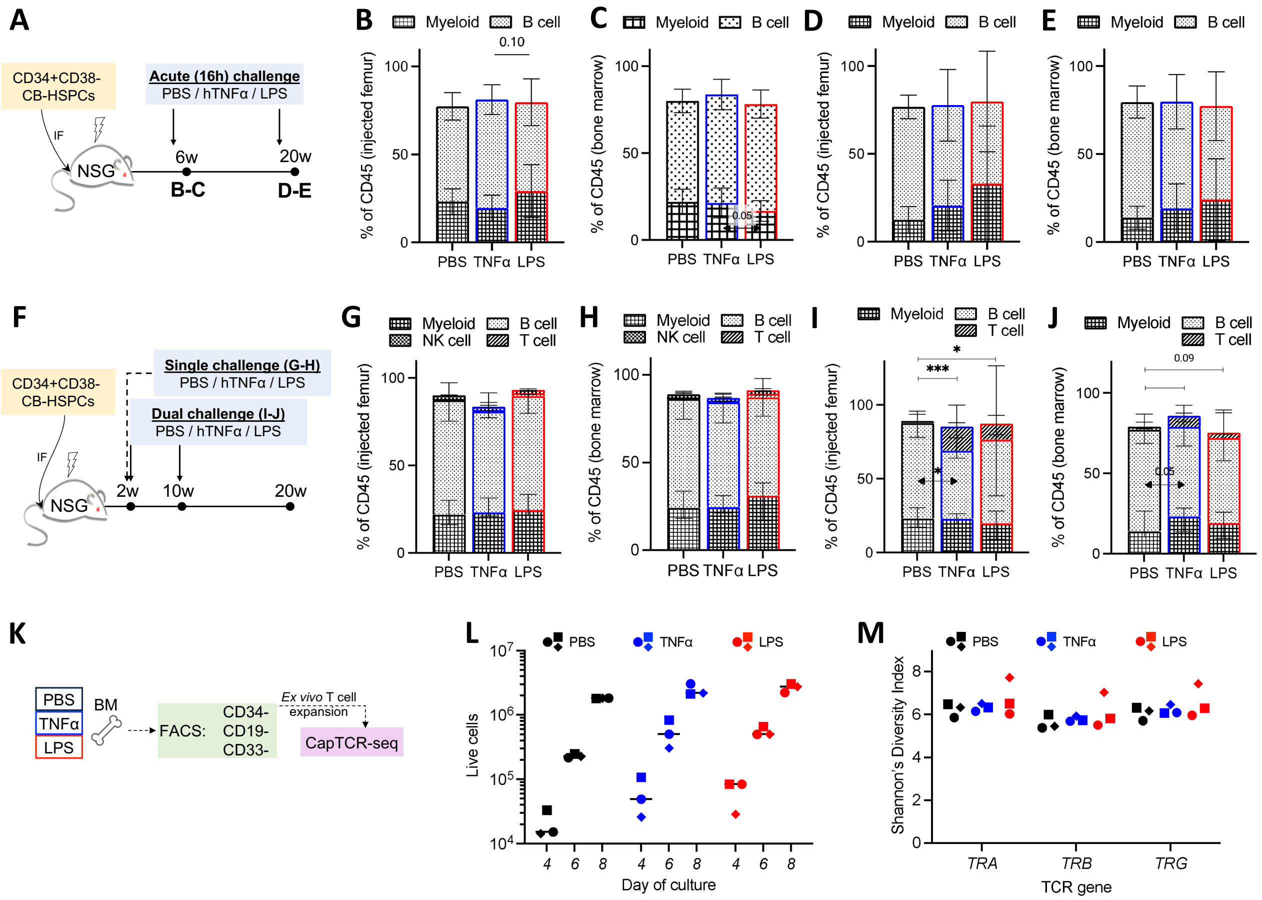


**Fig. S3. Lineage distribution in human HSC grafts after acute inflammatory challenge as well as after recovery.** **(A)** Experimental schematic for acute inflammatory challenge of xenografts with hTNFα or LPS at two time points. **(B-C)** Proportion of human B cell (CD19+) versus myeloid (CD33+) composition in injected or non-injected bone marrow of mice 16 hours after being subjected to inflammatory challenge at 6 weeks post-transplant. **(D-E)** Proportion of human B cell versus myeloid composition in injected or non-injected bone marrow of mice 16 hours after being subjected to inflammatory challenge at 20 weeks post-transplant. **(F)** Schematic for xenograft model of recovery from single (2 weeks only) or dual (2 weeks and 10 weeks) acute inflammatory challenge followed by recovery. **(G-H)** Proportion of human B cell versus myeloid versus T/NK cell (CD3+ and CD56+, respectively) composition in injected or non-injected bone marrow of mice at 20 weeks post-transplant; 18 weeks after being subjected to a single inflammatory challenge at 2 weeks post-transplant. **(I-J)** Proportion of human B cell versus myeloid versus T cell composition in injected or non-injected bone marrow of mice at 20 weeks post-transplant; 10 weeks after being subjected to two inflammatory challenges at 2 weeks, then 10 weeks post-transplant, respectively. Stacked bar plots for B/T/NK/Myeloid proportions represent median composition from the same experiments, and error bars reflect interquartile range. Data were compared using pairwise Mann-Whitney tests; *** $p<0.001$, ** $p<0.01$, * $p<0.05$, and $p<0.1$ is shown numerically, while $p>0.1$ is not shown. **(K-M)** T cells from dual challenge + 10w recovery xenografts (I-J) were expanded in culture and T cell receptor (TCR) repertoire was evaluated by CapTCR-seq (K). Live cell counts were determined via trypan blue exclusion (L) at the indicated time points. Shannon's diversity index was calculated for the unique sequences obtained for the three indicated TCR genes (M). Each point/symbol reflects data obtained from pooled T cells from a group of mice engrafted with different CB pools.

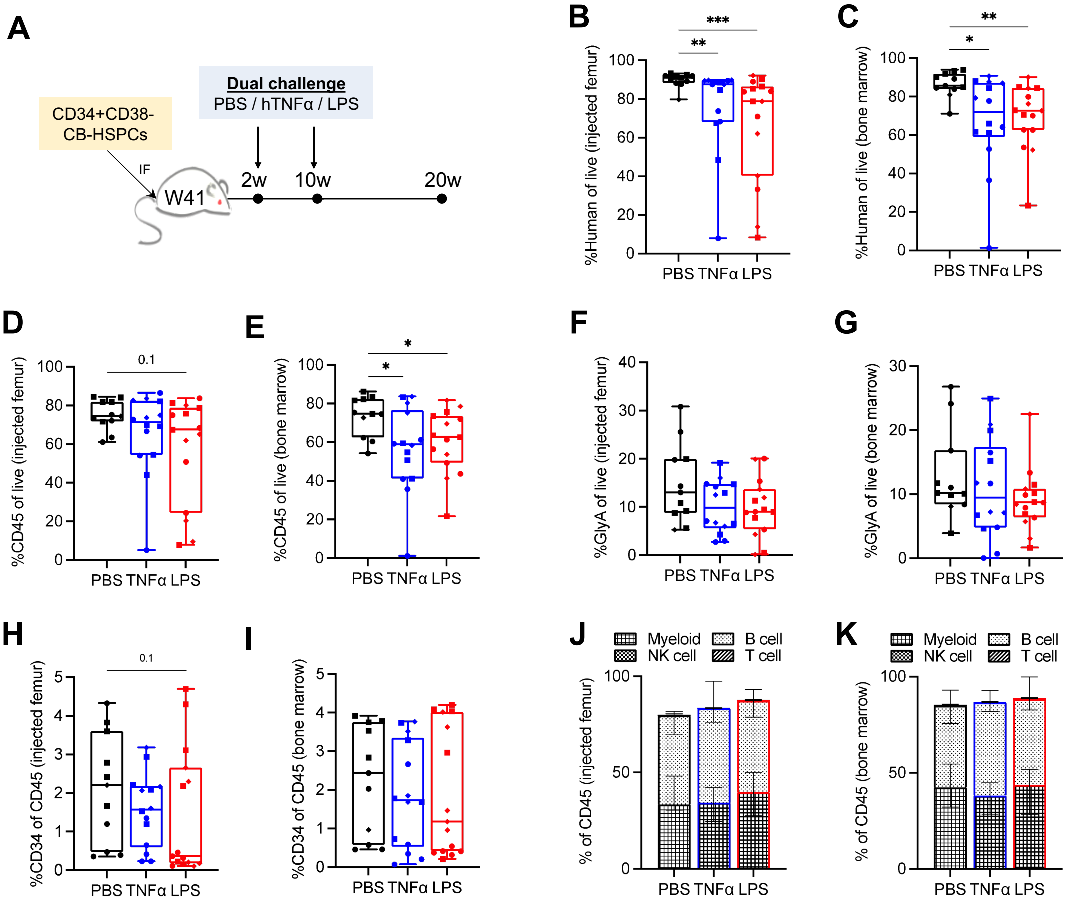


**Fig. S4. Recovery of human HSCs from inflammation in xenografted NSGW41 mice.**

**(A)** Schematic for xenograft model of recovery from dual (2 weeks and 10 weeks) acute inflammatory challenge in NSGW41 mice. **(B-G)** Engraftment of total human cells (B,C) - calculated by combining CD45+ (D,E) and GlyA+ (F,G) proportions - in injected and non-injected bone marrow from dual-challenged NSGW41 xenografts. **(H-K)** Proportion of CD34+CD19-CD33- progenitors (H,I), or lineage composition by myeloid (CD33+), B (CD19+), NK (CD56+) and T (CD3+) cells (J,K), in injected and non-injected bone marrow from dual-challenged NSGW41 xenografts. Engraftment data is presented as box plots with a line at the median, boxes showing quartiles and error bars showing range. Individual points show data for each animal used in the study (n=10-15), and different symbols reflect different CB pools (n=3). Stacked bar plots for B/T/NK/Myeloid proportions represent median composition from the same experiments, and error bars reflect interquartile range. Data were compared using pairwise Mann-Whitney tests; *** $p<0.001$, ** $p<0.01$, * $p<0.05$, and $p<0.1$ is shown numerically, while $p>0.1$ is not shown.

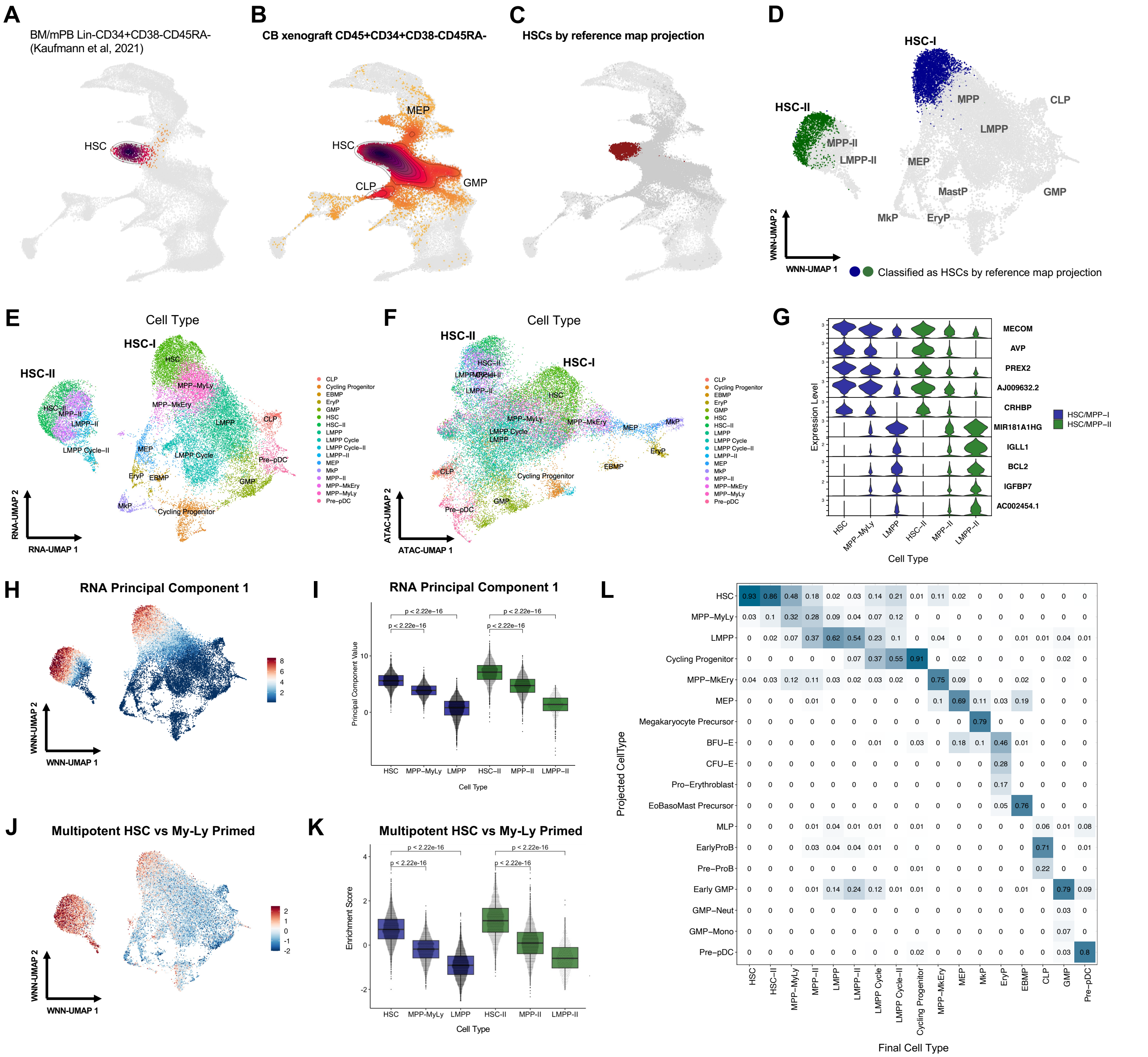


**Fig. S5. Cell type characterization of dual challenge CB xenograft scMultiome profiles.**

**(A)** BoneMarrowMap projection results of scRNA-seq from Lin-CD34+CD38-CD45RA- cells from Kaufmann et al 2021, sorted from bone marrow and mobilized peripheral blood samples. **(B)** Projection results of snRNA-seq from CD45+CD34+CD38-CD45RA- cells profiled by 10x Multiome, sorted from 20-week cord blood xenografts in the setting of dual inflammatory challenge and recovery. **(C)** 7,507 CB xenograft cells identified as HSCs by reference map projection from xenografted cells in (B). **(D)** UMAP of 27,492 xenografted HSPCs based on integrated RNA and ATAC embeddings from weighted nearest-neighbour (WNN) analysis. Cells that were classified as HSCs by reference map projection are colored; all other cells are shown in grey. **(E-F)** UMAP embeddings for xenografted HSPCs based on gene expression (E) and chromatin accessibility (F) modalities alone. **(G)** Normalized expression of marker genes specific to HSC or LMPP. **(H-I)** Value of gene expression principal component 1 (PC1) overlaid on the WNN UMAP embedding (H) and depicted as boxplots for HSC, MPP, LMPP, and HSC-II, MPP-II, LMPP-II populations (I). Statistical comparisons are made with a Wilcoxon rank-sum test. **(J-K)** Normalized enrichment score (AUCell) of a signature of multipotent human HSC compared to myelo-lymphoid lineage restricted human HSC from Belluschi *et al* 2018. Enrichment is shown on the WNN UMAP embedding (J) and depicted as boxplots for HSC, MPP, LMPP, and HSC-II, MPP-II, LMPP-II populations (K). Statistical comparisons are made with a Wilcoxon rank-sum test. **(L)**

Confusion matrix depicting the final cell types assigned by focused RNA + ATAC based clustering (columns) with projected cell types based on reference gene expression profiles from BoneMarrowMap (rows). Shown proportions are normalized by column.

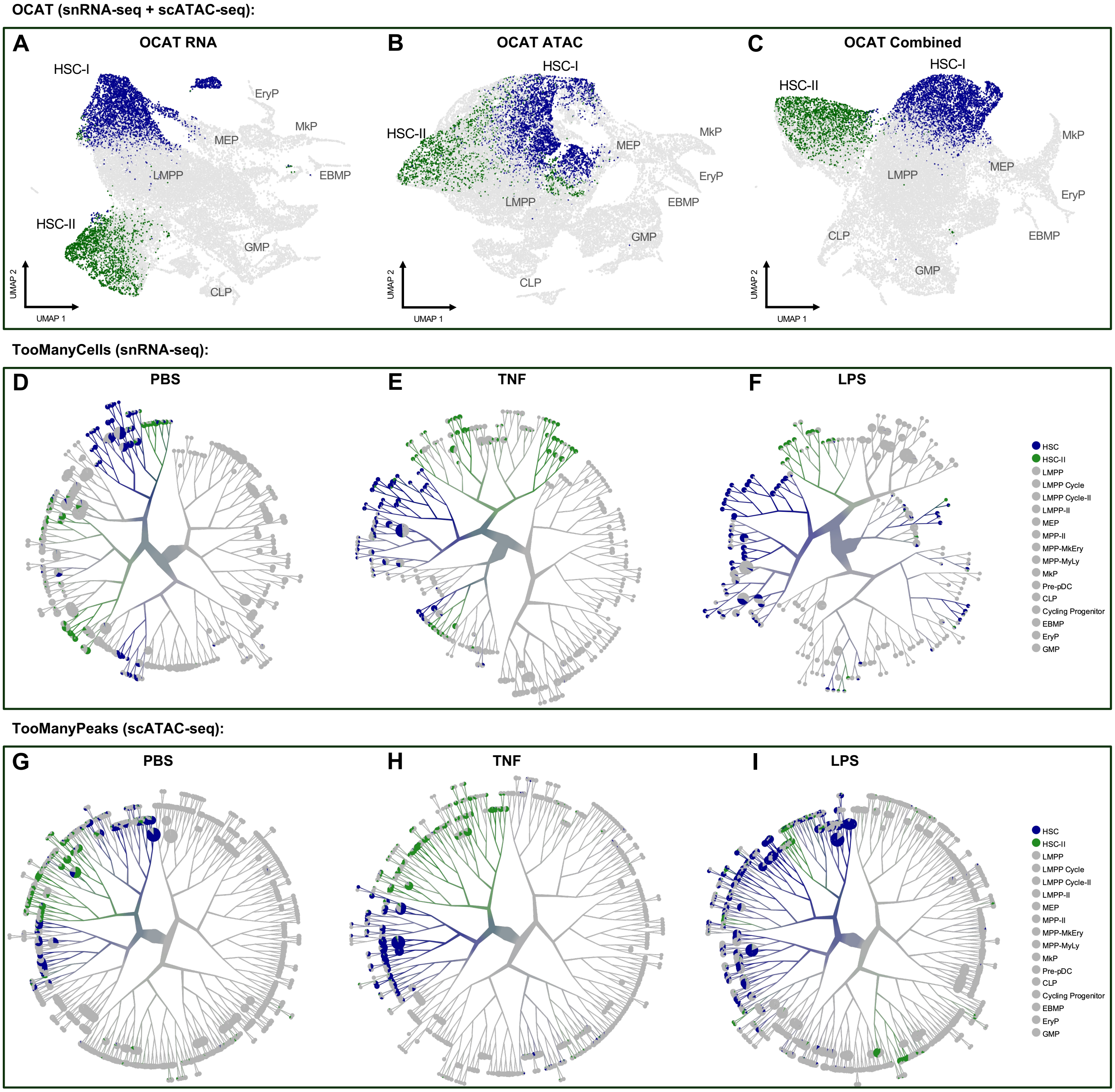


**Fig. S6. Alternative approaches to neighborhood analysis of HSC-I and HSC-II.**

**(A-C)** Alternative reduced dimensional embeddings through One Cell At a Time (OCAT), which constructs "ghost" cells based on neighborhood centroids and generates a sparse encoding for each cell in the dataset based on its proximity to each "ghost" cell, prior to downstream UMAP construction. These embeddings were generated on the basis of gene expression (A), chromatin accessibility (B), and combined gene expression + chromatin accessibility (C). HSC-I and HSC-II are colored in blue and green, respectively, within each embedding. **(D-F)** Cell clade visualization from gene expression data by TooManyCells, shown for cells from each treatment condition. Clades containing HSC-I and HSC-II are colored in blue and green, respectively. **(G-I)** Cell clade visualization from chromatin peak accessibility by TooManyPeaks, shown for cells from each treatment condition. In this analysis, the input is the chromatin peak matrix called by MACS2 also used as an input for classical scATAC analysis by Signac. Clades containing HSC-I and HSC-II are colored in blue and green, respectively.

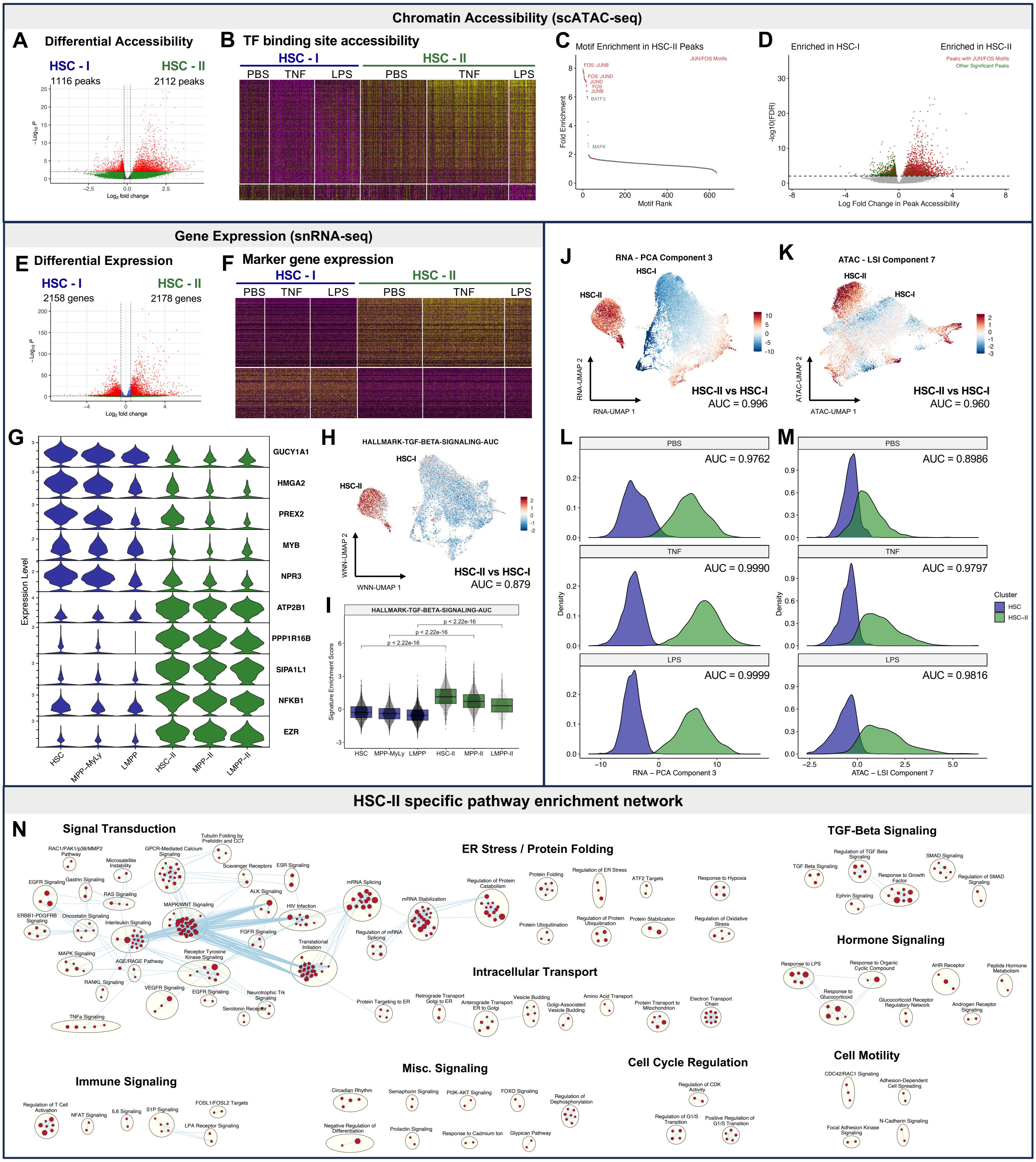


**Fig. S7. Epigenetic and transcriptional characterization of HSC-I and HSC-II.**

**(A)** Differentially accessible (DA) peaks between HSC-II and HSC-I, performed by creating pseudo-bulks by donor sex and treatment condition. Significant peaks were called at an FDR < 0.05. **(B)** Heatmap of DA transcription factor binding sites (TFBS) from ReMap2020; normalized chromVAR enrichment scores are shown. **(C)** Fold enrichment of TF motifs, from the JASPAR database, within DA peaks specific to HSC-II. **(D)** Presence of JUN-FOS motifs within DA peaks between HSC-II and HSC-I. **(E-F)** Differentially expressed (DE) genes between HSC-II and HSC-I, identified through pseudo-bulk analysis as per (A). **(G)** Normalized expression of top HSC-I and HSC-II markers within HSC, MPP, LMPP subsets. **(H-I)** Normalized enrichment score of the TGF-β signaling pathway on the WNN

UMAP embedding (H) and across HSC, MPP, LMPP subsets (I). Statistical comparisons are made with a Wilcoxon rank-sum test. **(J-M)** Value of reduced dimensions gene expression PC 3 (J, L) and chromatin accessibility LSI component 7 (K, M) on the WNN UMAP embedding, and separability across HSC-I and HSC-II, stratified by treatment condition. **(N)** Enrichment map of GO biological pathways enriched in HSC-II by GSEA based on differential expression analysis from (E) adjusting for treatment condition as a covariate. Each pathway (red dot) is clustered based on overlap with other pathways. Only pathways enriched at FDR < 0.01 are shown, singleton pathways were removed. There were no biological pathways uniquely enriched in HSC-I at FDR < 0.01.

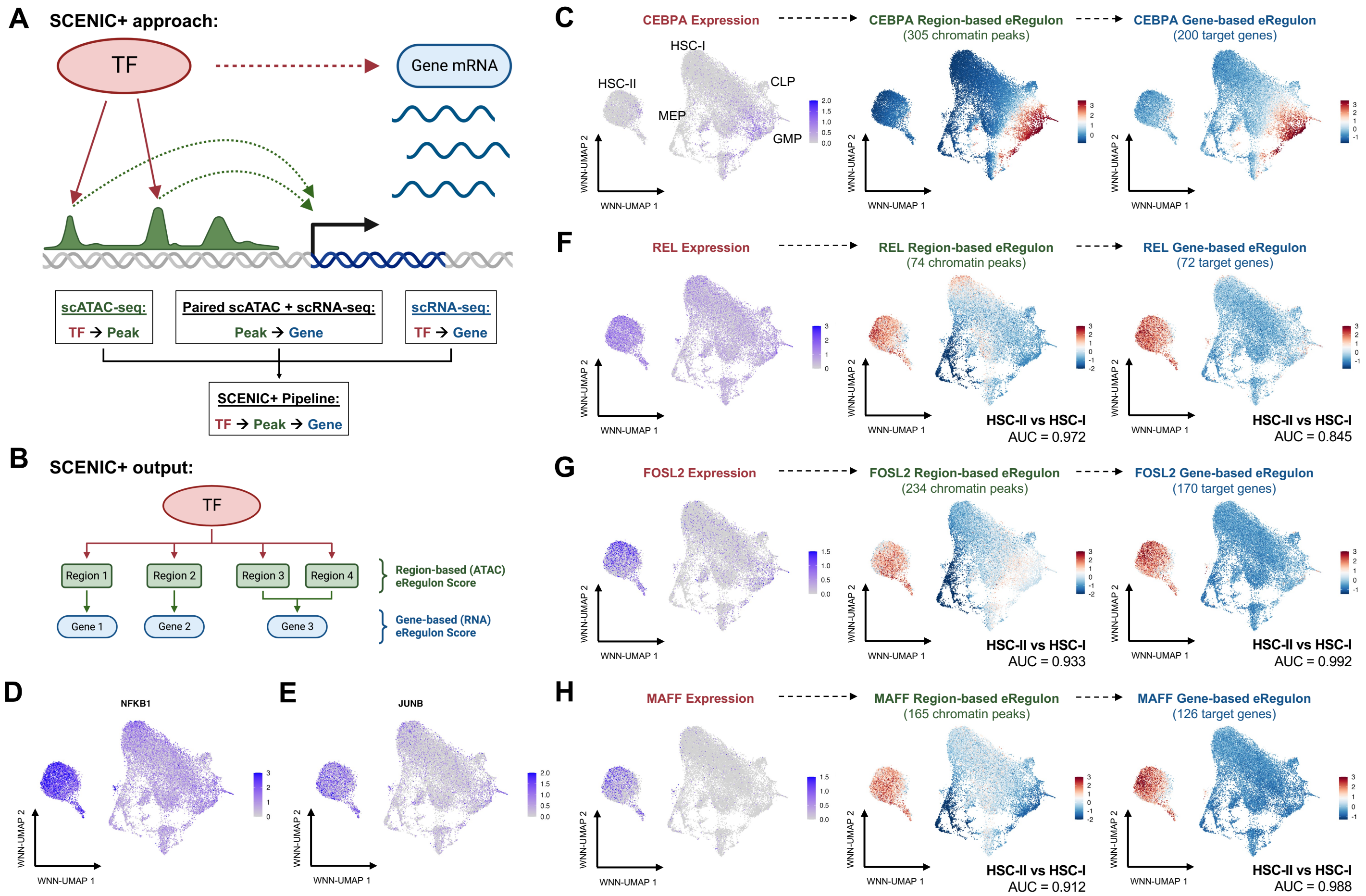


**Fig. S8. Multiomic inference of gene regulation using SCENIC+.**

**(A)** Schematic outlining multi-modal inference of gene regulation by SCENIC+. Transcription factor (TF) to target gene relationships are inferred through SCENIC analysis on single-cell transcriptome data. TF to chromatin peak relationships are inferred through single-cell chromatin accessibility data. Finally, cis-regulation between chromatin peaks and nearby genes is inferred through paired chromatin accessibility and gene expression profiled in the same cells. These three layers of information are integrated together to infer regulation from TF to chromatin peak to target gene expression. **(B)** Schematic illustrating SCENIC+ output in estimating activity, or "eRegulon score", of an individual TF based on accessibility of target chromatin regions and expression of target genes regulated through those regions. **(C)** Validation of SCENIC+ with CEBPA, a known master regulator of myeloid fate. CEBPA transcript expression, chromatin region-based eRegulon score, and gene expression-based eRegulon score are shown. **(D-E)** Normalized expression of NFKB1 (C) and JUNB (D) transcripts on the WNN UMAP embedding. **(F-H)** Depiction of TF expression, region-based eRegulon score, and gene-based eRegulon scores of key HSC-II specific regulators including REL (F), FOSL2 (G), and MAFF (H).

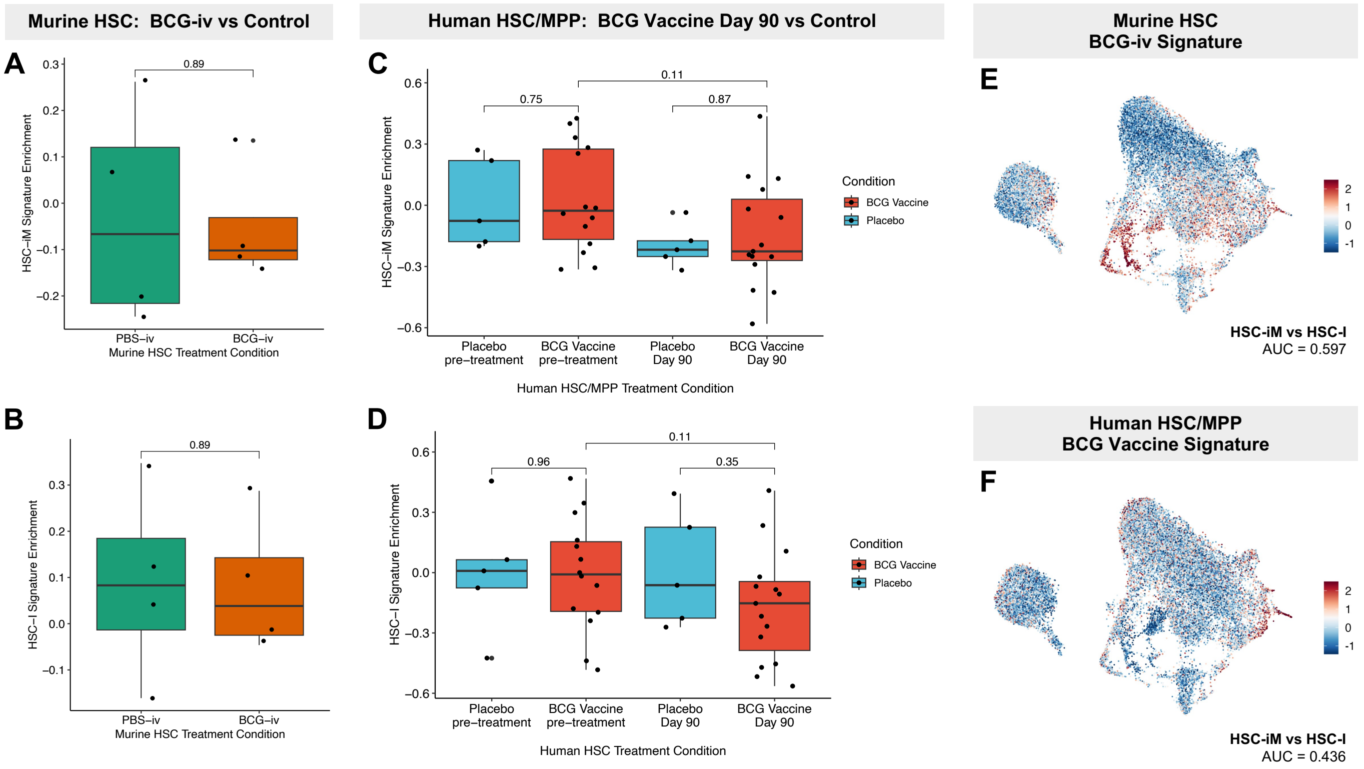


**Fig. S9. Inflammatory memory in HSCs is distinct from memory of prior BCG vaccination.**

**(A-B)** Enrichment of HSC-iM vs HSC-I marker genes within murine HSCs (LSK+CD150+) treated with intravenous infusion of either the BCG vaccine or PBS. Enrichment scores were calculated in bulk RNA-seq data by GSVA. Human marker gene names were converted to murine orthologs using the babelgene R package. (A) Enrichment of top 200 genes specific to HSC-iM compared to HSC-I. (B) Enrichment of top 200 genes specific to HSC-I compared to HSC-iM. **(C-D)** Enrichment of HSC-iM vs HSC-I marker genes within HSC/MPPs (CD34+CD38-CD45RA-) from human volunteers receiving either a BCG vaccine or placebo. Samples were collected either pre-vaccination or at day 90 following vaccination. Enrichment scores were calculated in bulk RNA-seq data by GSVA. (C) Enrichment of top 200 genes specific to HSC-iM compared to HSC-I. (D) Enrichment of top 200 genes specific to HSC-I compared to HSC-iM. **(E-F)** Enrichment of BCG-specific marker genes within the xenograft multi-ome dataset. Enrichment scores were calculated in scRNA-seq data by AUCell. (E) Enrichment of a murine HSC BCG-iv signature comprised of 490 genes enriched in BCG-iv HSCs at FDR < 0.05 and LFC > 1, which were converted to human orthologs using the babelgene R package. (F) Enrichment of an HSC/MPP BCG vaccination signature comprised of 206 genes enriched in HSC/MPPs at Day 90 after BCG vaccination compared to HSC/MPPs prior to vaccination at FDR < 0.05 and LFC > 1.

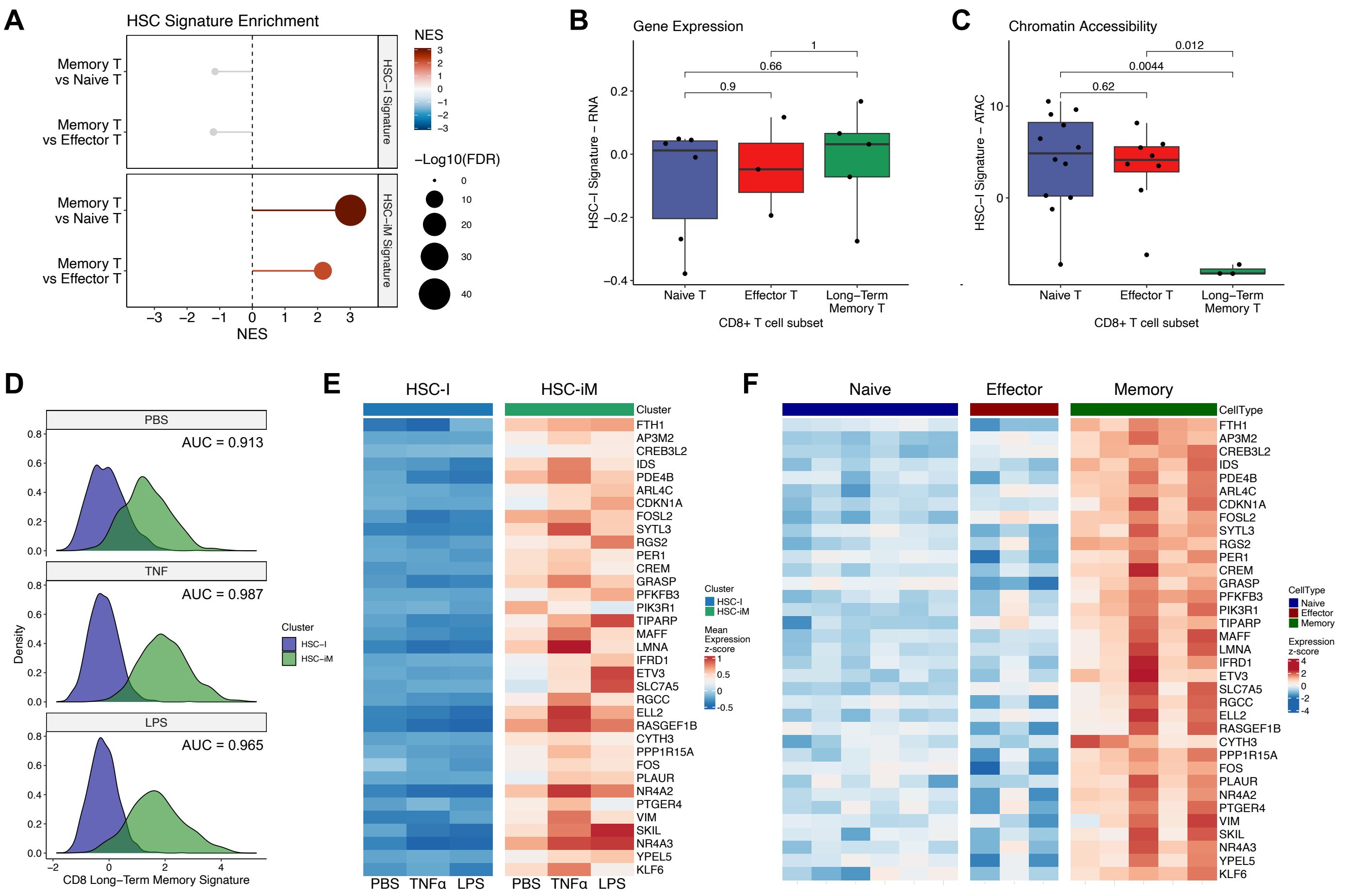


**Fig. S10. Conserved gene expression programs between HSC-iM and functional T cell memory.**

**(A)** GSEA results depicting enrichment of HSC-I and HSC-iM specific gene expression signatures between human T cell subsets. **(B)** Enrichment of an HSC-I gene expression signature, comprised of the top 200 differentially expressed genes specific to HSC-I (vs HSC-iM), within human T cell subsets profiled by RNA-seq. **(C)** Enrichment of an HSC-I chromatin accessibility signature, comprised of 2,475 differentially accessible regions specific to HSC-I (vs HSC-iM) within human T cell subsets profiled by ATAC-seq. **(D)** Enrichment of a long-term T cell memory signature within HSC-iM and HSC-I, stratified by treatment condition. **(E-F)** Expression of 35 genes shared between the 200 gene HSC inflammatory memory signature and 257 gene T cell memory signature, (E) within HSC-I and HSC-II in each treatment condition and (F) within human T cell subsets.

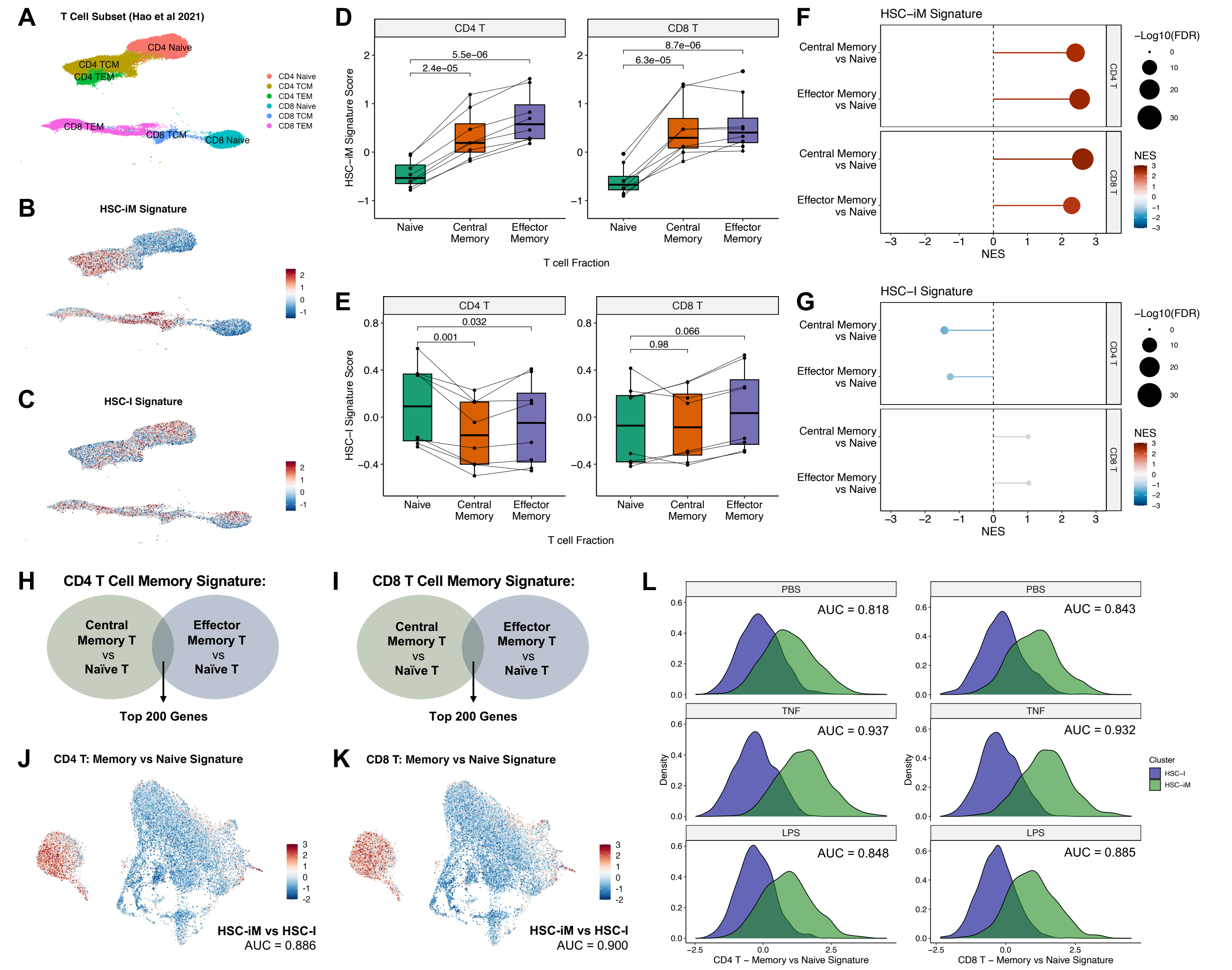


**Fig. S11. HSC Inflammatory Memory signature in CD4 and CD8 Memory T cells.**

**(A)** Integrated UMAP embedding of human PBMC CITE-seq from Hao *et al.,* 2021, incorporating gene expression together with immunophenotype. Only Naïve, Central Memory T (TCM), and Effector Memory T (TEM) CD4 and CD8 T cells are shown. **(B-C)** HSC signature enrichment in human T cell data, represented by the scaled AUCell enrichment score. These are shown for (B) top 200 HSC-iM specific genes or (C) top 200 HSC-I specific genes. **(D-E)** HSC signature enrichment in human T cell subsets at the level of individual donors from Hao *et al* 2021. Scaled AUCell enrichment scores were averaged across cells in each subset within each of 8 human donors, subsets from each donor are connected through by a line. Comparisons were performed through a paired t-test. These are shown for (D) top 200 HSC-iM specific genes or (E) top 200 HSC-I specific genes. **(F-G)** GSEA results depicting HSC signature enrichment in differential expression results between memory and naïve T cells subsets from Hao *et al* 2021, utilizing pseudobulk profiles and adjusting for donor ID as a covariate. Positive enrichment indicates memory T cell subsets compared to naïve subsets. Results are faceted by CD4 and CD8 T cell type. These are shown for (F) top 200 HSC-iM specific genes or (G) top 200 HSC-I specific genes. **(H-I)** Derivation of Memory vs Naïve subset signature within (H) CD4 T cells and (I) CD8 T cells. Among genes enriched in both TCM vs Naïve and TEM vs Naïve comparisons at LFC > 1 and FDR < 0.01, test statistics were averaged between comparisons and the top 200 genes were select by mean test statistic value. **(J-K)** Enrichment of memory T cell marker genes from (J) CD4 T cells and (K) CD8 T cells within the xenograft multi-ome dataset. Enrichment scores were calculated in scRNA-seq data by AUCell. **(L)** Enrichment of memory T cell marker genes within HSC-iM and HSC-I, stratified by treatment condition.

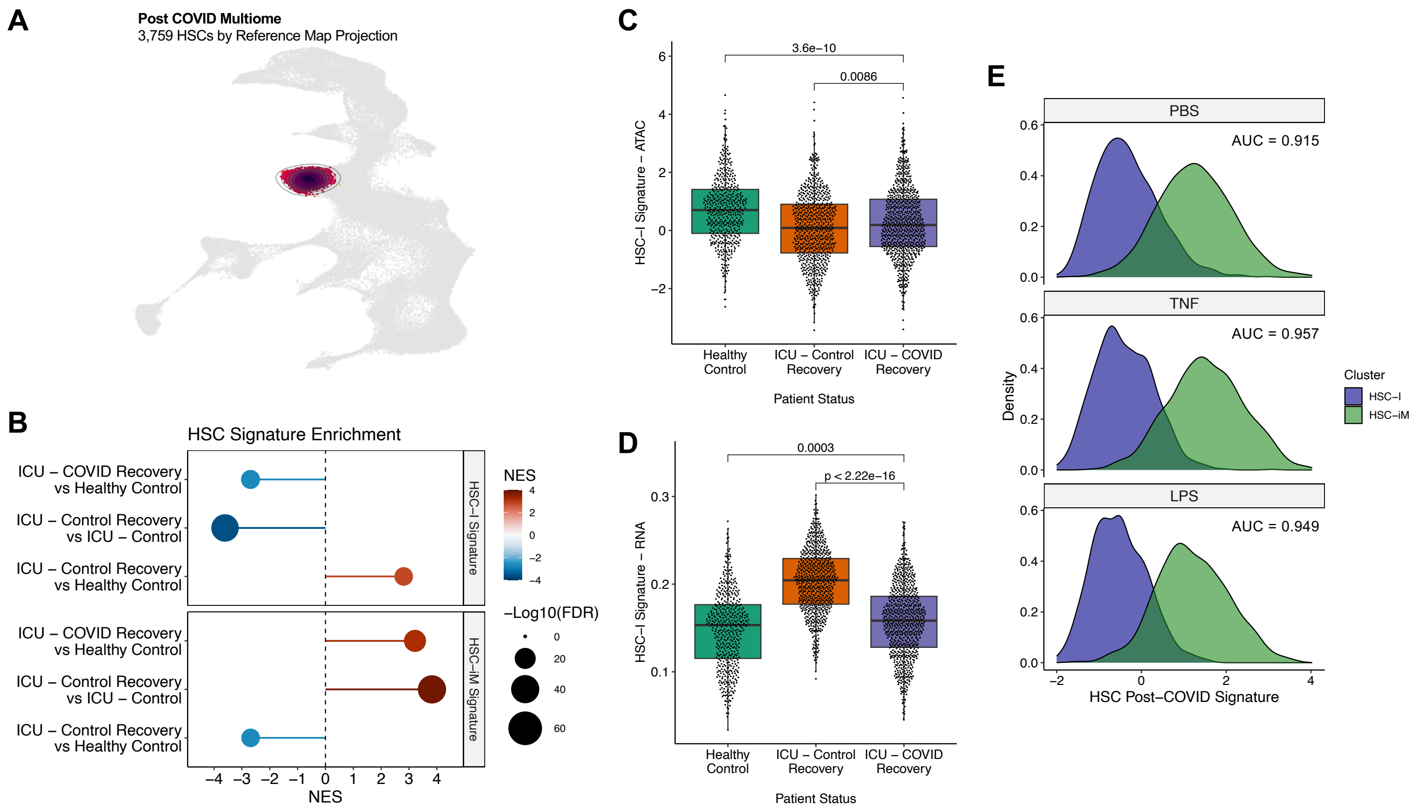


**Fig. S12. HSC Inflammatory Memory signature in HSCs following severe COVID-19 illness.**

**(A)** 3,759 cells from Cheong *et al* 2023 classified as HSCs through reference map projection. **(B)** GSEA results depicting enrichment of HSC-I and HSC-iM specific gene expression signatures between HSCs across conditions spanning Healthy Control, ICU Control, and Severe COVID-19 Recovery. **(C-D)** Enrichment of HSC-I specific signatures across conditions, scored on the basis of chromatin accessibility (C) and gene expression (D). Statistical comparisons were made using Wilcoxon rank-sum tests. **(E)** Enrichment of a post-COVID HSC gene expression signature between HSC-I and HSC-iM within the xenograft scMultiome data, stratified by treatment condition.

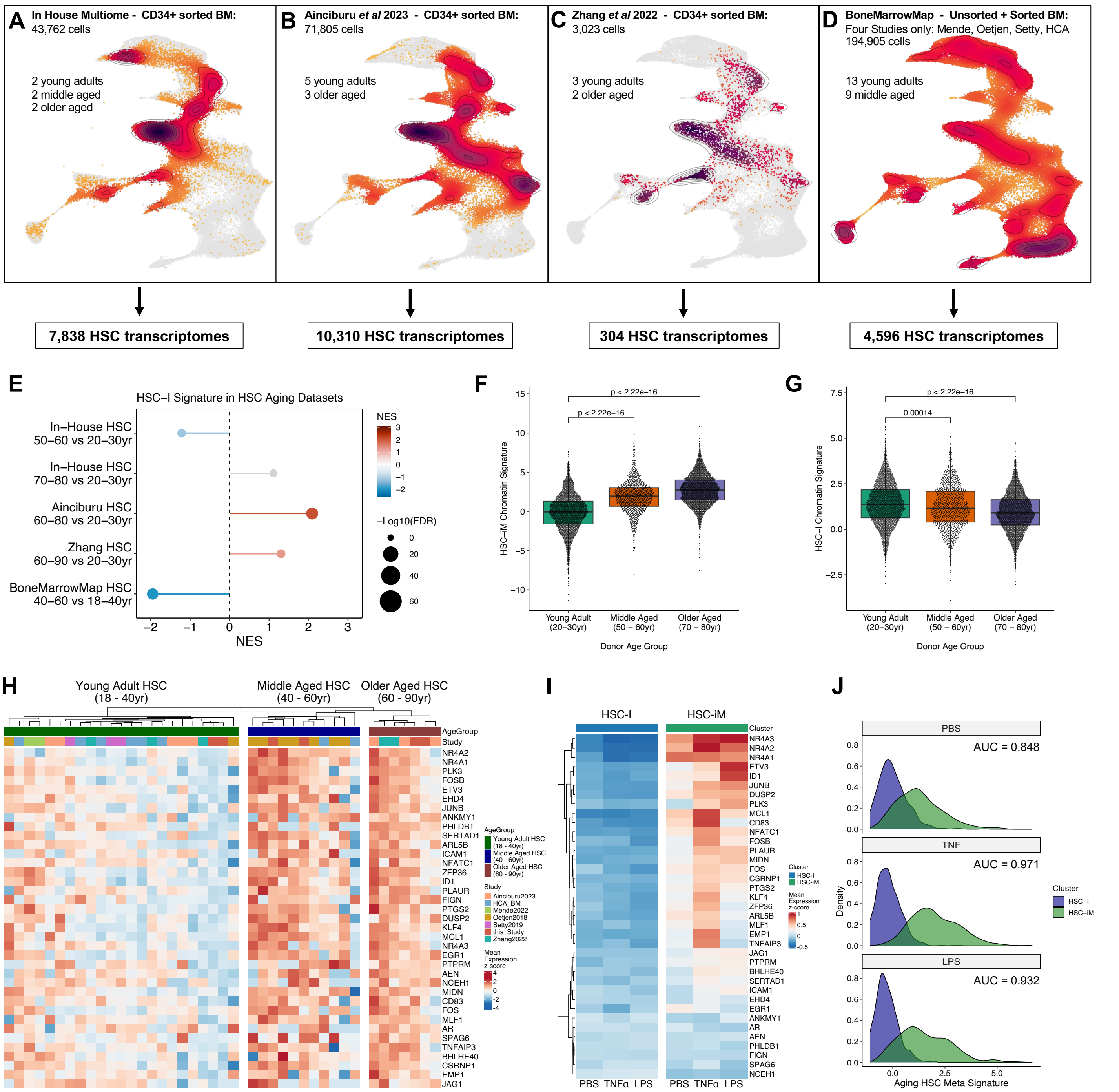


**Fig. S13. HSC Inflammatory Memory signature in HSCs across a human lifetime.**

**(A-D)** *In silico* purification of HSCs by BoneMarrowMap reference map projection from single-cell transcriptome datasets of human HSPCs from young adult (YA), middle aged (MA), and older aged (OA) bone marrow aspirates. (A) 10x scMultiome snRNA-seq + scATAC-seq from 43,762 CD34+ HSPCs from 6 donors (2 YA, 2 MA, 2 OA) sequenced in house, yielding 7,838 HSCs. (B) 10x v2 and v3 scRNA-seq from 71,805 CD34+ HSPCs from 8 donors (5 YA, 3 OA) profiled in Ainciburu *et al* 2023, yielding 10,310 HSCs. (C) STRT-seq from 3,023 CD34+ HSPCs from 5 donors (3 YA, 2 OA) profiled in Zhang *et al* 2022, yielding 304 HSCs. (D) 10x v2 scRNA-seq from four datasets within BoneMarrowMap: Oetjen *et al* 2018 (unsorted BM), Human Cell Atlas BM (unsorted BM), Setty *et al* 2019 (CD34+ BM), and Mende *et al* 2022 (CD34+ BM). Among 22 donors with ≥10 HSCs (13 YA, 9 MA), 4,596 HSCs were retained. **(E)** GSEA results depicting HSC-I gene expression signature enrichment in differential expression results between five separate comparisons of older aged (OA) and middle aged (MA) vs young adult (YA) human HSC. **(F-G)** ChromVAR enrichment scores of accessible chromatin peaks specific to HSC-iM (F) and HSC-I (G) subsets in scATAC-seq from YA, MA, and OA HSCs within the in-house scMultiome dataset. **(H)** Expression of 37 shared genes commonly upregulated in aged HSCs through meta-analysis of independent HSC aging datasets in (A-D). Each column is a pseudo-bulk profile of HSCs within an individual donor from each aging study. **(I)** Expression of 37 shared HSC aging genes within HSC-I and HSC-II in each treatment condition. **(J)** Enrichment score of 37-gene HSC aging meta-signature between HSC-I and HSC-iM, stratified by treatment condition.

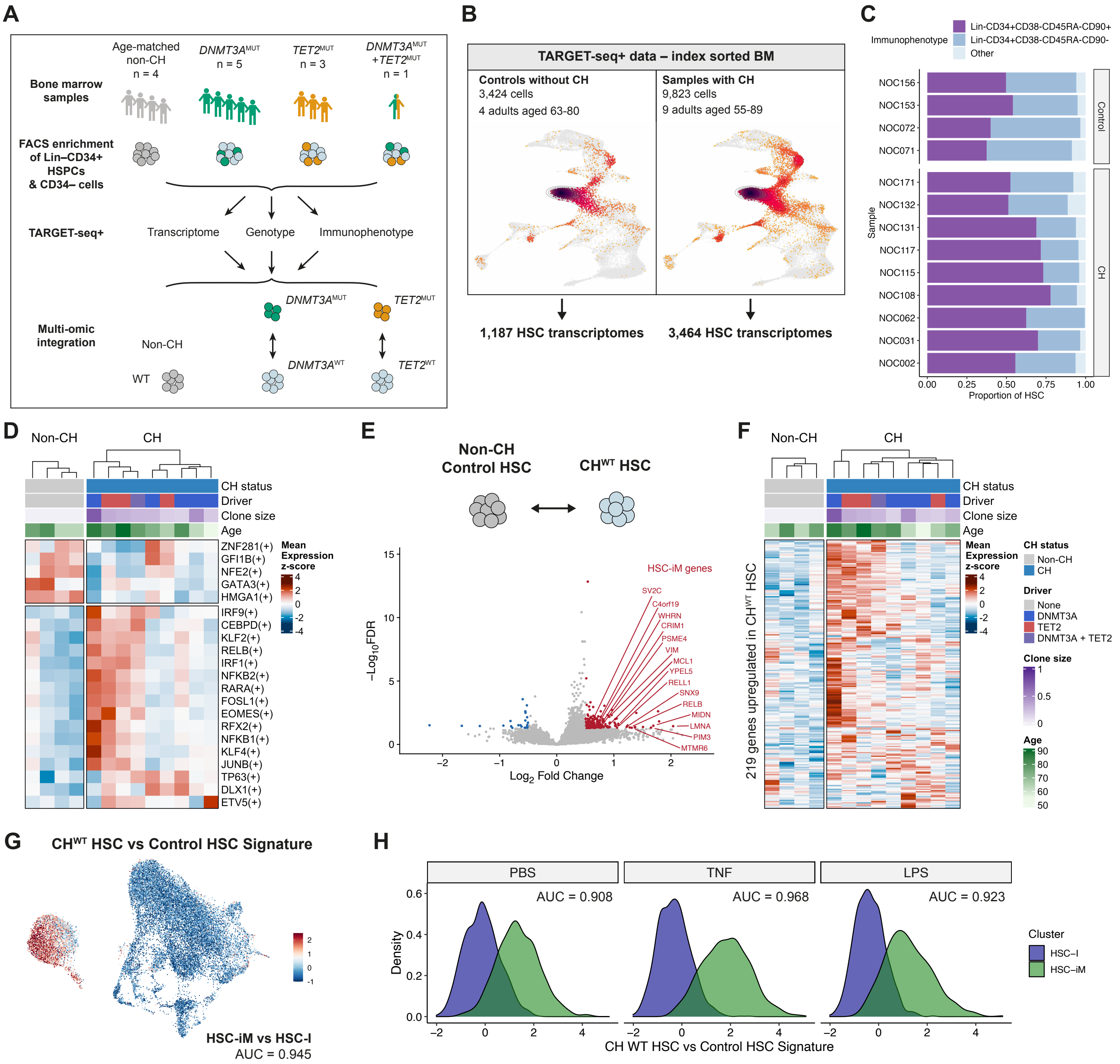


**Fig. S14. Inflammatory Memory Program in Human Clonal Hematopoiesis.**

**(A)** Schematic outlining TARGET-seq+ analysis of BM samples from donors with CH and age-matched controls without CH (non-CH). Samples were FACS enriched for Lin$^-$CD34$^+$ HSPCs, Lin$^-$CD34$^+$CD38$^-$ cells and CD34$^-$ cells and processed using TARGET-seq+ obtaining transcriptome, targeted genotyping, and flow cytometry index immunophenotyping for each cell. **(B)** *In silico* purification of HSCs by BoneMarrowMap reference map projection from the single-cell TARGET-seq+ dataset of index-sorted human bone marrow samples, which were enriched for CD34+ HSPCs. Left: 3,424 cells from 4 donors without CH mutations (aged 63-80), yielding 1,187 HSCs. Right: 9,823 cells from 9 donors with *DNMT3A* and *TET2* CH mutations (aged 55-89), yielding 3,464 HSCs. **(C)** Immunophenotypic profiles of HSCs purified by BoneMarrowMap reference map projection. **(D)** Expression of regulons differentially expressed (with FDR < 0.1) between CH$^{WT}$ HSCs and HSCs from non-CH donors, performed by linear mixed model adjusting for donor ID, donor age, donor sex, and FACS sort batch. Each column represents a pseudo-bulk from one donor. Donors are annotated with their CH status, driver mutation, the clone size in bulk BM sequencing and their age. **(E)** Differentially expressed genes (DEGs) between CH$^{WT}$ HSCs from 9 donors and control HSCs from 4 non-CH donors, performed by dream linear mixed model adjusting for donor ID, donor age, donor sex, and FACS sort batch. Significant genes were called with FDR < 0.05 and log2FC > 0.5. DEGs in the HSC-iM signature are labelled. **(F)** Expression of 219 genes upregulated in $CH^{WT}$ HSCs from (E) within $CH^{WT}$ HSCs and HSCs from non-CH donors. Donors are annotated as per (D). **(G)** Enrichment of the $CH^{WT}$ HSC versus Control HSC signature from (E) within the xenograft multi-ome dataset. Enrichment scores were calculated in scRNA-seq data by AUCell. **(H)** Enrichment of the $CH^{WT}$ HSC versus Control HSC signature within HSC-iM and HSC-I, stratified by treatment condition.

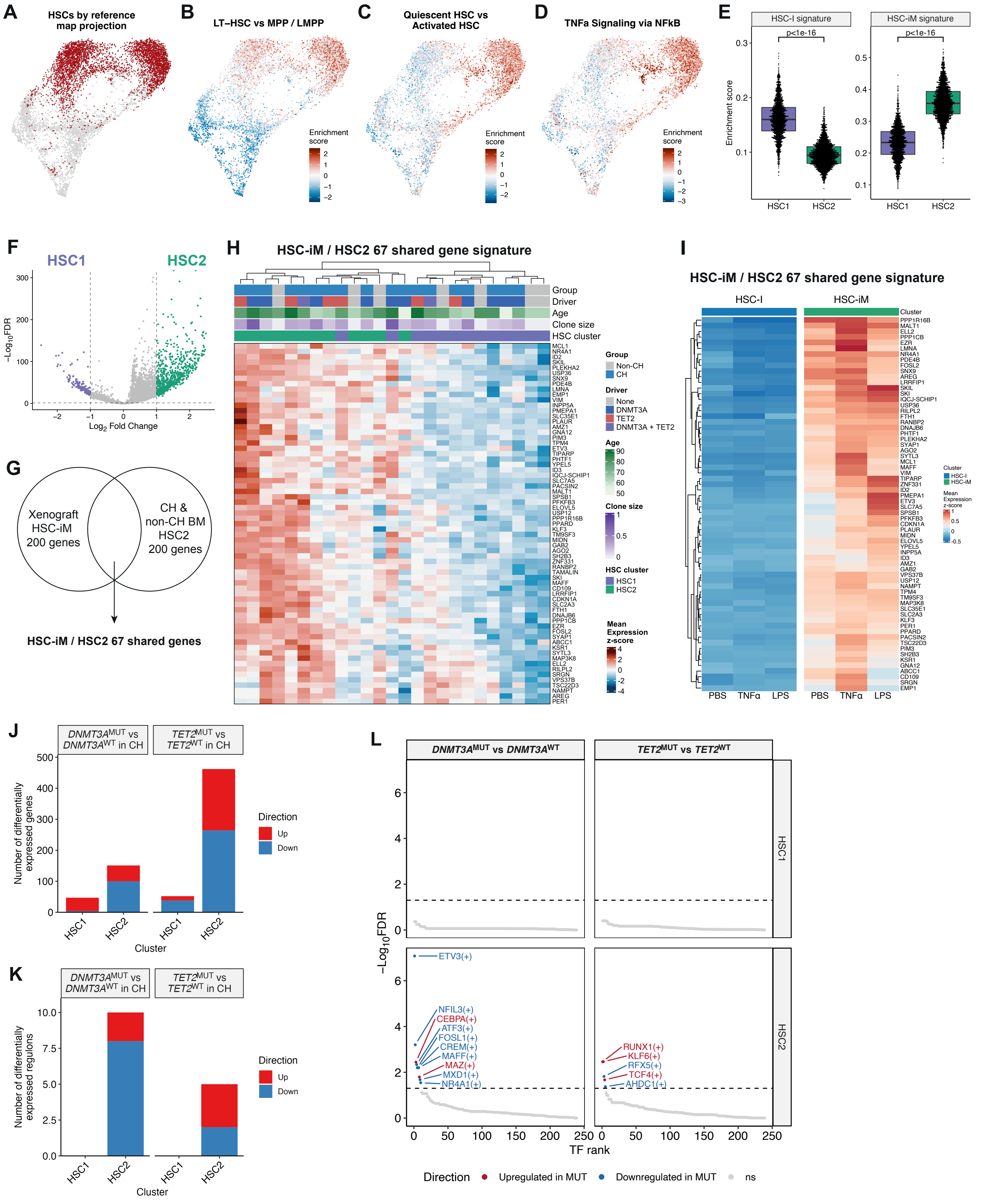


**Fig. S15. Identification of an Inflammatory Memory HSC Population in Human Clonal Hematopoiesis.**

**(A)** UMAP of 8,059 bone marrow HSPCs from 9 donors with CH and 4 age-matched samples without CH, after feature weight derivation with the Self-Assembling Manifolds (SAM) algorithm. The 4,651 cells identified as HSCs by reference map projection are highlighted. **(B-D)** Normalized signature enrichment scores (AUCell), overlaid on the SAM UMAP embedding. (B) Enrichment of an LT-HSC-specific gene signature from sorted fraction bulk RNA-seq in CB. (C) Enrichment of a gene expression signature specific to uncultured, quiescent, CB LT-HSC relative to CB LT-HSC activated from 96hr in *in vitro* culture. (D) Enrichment of the Hallmark TNFα signaling via NF-kB signature. **(E)** Enrichment of the HSC inflammatory memory signature within the HSC1 and HSC2 populations identified in the TARGET-seq+ dataset of CH and non-CH donors. Statistical comparisons were made using Wilcoxon rank-sum tests. **(F)** Differentially expressed genes (DEGs) between HSC1 and HSC2 populations identified in CH and non-CH donors, performed by dream linear mixed model adjusting for donor sex. Significant genes were called with FDR < 0.05 and log2FC > 1. **(G)** Intersection of the 200 gene HSC inflammatory memory signature and the top 200 genes upregulated in HSC2 compared to HSC1 from (F) identified 67 shared genes. **(H)** Expression of the 67 genes shared between the HSC-iM and HSC2 signatures. Each column is a pseudo-bulk profile within an individual donor from each HSC cluster. Donors are annotated with their CH status, driver mutation, the clone size in bulk BM sequencing, their age, and the HSC cluster. **(I)** Expression of the 67 marker genes shared between HSC-iM and HSC2 within HSC-I and HSC-iM in each treatment condition. **(J)** Number of differentially expressed genes between $CH^{MUT}$ and $CH^{WT}$ cells within the HSC1 and HSC2 clusters. Significant genes were called with FDR < 0.1. **(K)** Number of differentially expressed regulons between $CH^{MUT}$ and $CH^{WT}$ cells within the HSC1 and HSC2 clusters. TF activity was inferred by SCENIC. **(L)** Differential TF activity between $CH^{MUT}$ and $CH^{WT}$ cells in either the HSC1 or HSC2 cluster. Y-axis portrays the significance level of differential enrichment by a linear mixed model accounting for donor identity. TFs are colored based on significance at FDR < 0.05.
